# Dimension reduction and denoising of single-cell RNA sequencing data in the presence of observed confounding variables

**DOI:** 10.1101/2020.08.03.234765

**Authors:** Mo Huang, Zhaojun Zhang, Nancy R. Zhang

## Abstract

Confounding variation, such as batch effects, are a pervasive issue in single-cell RNA sequencing experiments. While methods exist for aligning cells across batches, it is yet unclear how to correct for other types of confounding variation which may be observed at the subject level, such as age and sex, and at the cell level, such as library size and other measures of cell quality. On the specific problem of batch alignment, many questions still persist despite recent advances: Existing methods can effectively align batches in low-dimensional representations of cells, yet their effectiveness in aligning the original gene expression matrices is unclear. Nor is it clear how batch correction can be performed alongside data denoising, the former treating technical biases due to experimental stratification while the latter treating technical variation due inherently to the random sampling that occurs during library construction and sequencing. Here, we propose SAVERCAT, a method for dimension reduction and denoising of single-cell gene expression data that can flexibly adjust for arbitrary observed covariates. We benchmark SAVERCAT against existing single-cell batch correction methods and show that while it matches the best of the field in low-dimensional cell alignment, it significantly improves upon existing methods on the task of batch correction in the high-dimensional expression matrix. We also demonstrate the ability of SAVERCAT to effectively integrate batch correction and denoising through a data down-sampling experiment. Finally, we apply SAVERCAT to a single cell study of Alzheimer’s disease where batch is confounded with the contrast of interest, and demonstrate how adjusting for covariates other than batch allows for more interpretable analysis.

## Introduction

The identification of and adjustment for observed confounding variables has been extensively studied in the context of high-throughput sequencing experiments. An important category of such confounding variables is batch effects. Leek et al. [1] define batch effects to be subgroups of measurements which occur in different settings and result in unwanted variation in the measured data. For example, samples sequenced in different rounds can exhibit strong batch effects. In single-cell RNA sequencing, batch effects have been shown to introduce large unwanted variation in the data matrix [2, 3]. A large number of methods have been developed to remove batch effects [4-10]. Most methods focus on aligning the low-dimensional representations of cells across batches, while a few also output a batch-adjusted gene expression matrix. Accurate correction for batch effects in the gene expression matrix, and not just in its low-dimensional representation, is important for analyses such as differential expression and gene-pair correlation analysis. However, there has been limited evaluation on the effectiveness of existing methods to adjust for batch in the high dimensional gene expression matrix.

It is also well known that batch is not the only variable that can cause unwanted variation in scRNA-seq experiments. As scRNA-seq is now ubiquitously applied to human patient samples [11-13] and large cohorts in population-wide expression quantitative trait locus (eQTL) studies [14-16], subject-level variables such as age, sex, and ancestry may confound the question of interest. On top of these subject level covariates are the technical differences between samples in their preparation and processing time, to which batch effects are usually attributed [17-19]. In addition, cell-level covariates such as library size, cell cycle, and mitochondrial RNA content, the latter often used as a proxy for cell quality [20], may need to be accounted for in a proper analysis [21-23]. Thus, one can view the batch effect correction problem within the more general framework of observed covariate adjustment and modeling.

Separate from the batch correction literature for scRNA-seq, sophisticated denoising methods have been developed to increase the signal-to-noise ratio of single cell expression data [24-28]. The goal of denoising is to recover a gene expression matrix that mimics what we would have observed if we sequenced the cDNA libraries at higher depth. Thus, while covariate adjustment removes unwanted variation in the data that are due to observed sources, denoising reduces technical variation that are due to the random sampling of molecules during library preparation and sequencing. Most denoising methods assume that the true gene expression matrix can be approximated by a low-dimensional manifold, the recovery of which by representation learning is an active area of research [29-31]. Existing representation learning and denoising methods do not allow for the adjustment of general observed covariates, and thus, in current analysis pipelines, it is unclear how to integrate the batch correction and denoising steps.

Here we present a method called SAVERCAT for manifold learning, dimension reduction, and denoising of single-cell RNA-seq data that can flexibly adjust for observed covariates at both the subject and cell level. SAVERCAT is based on an adaptation of the conditional variational autoencoder (CVAE) [32] with a modified objective function which encourages disentanglement of latent and observed factors. First, the performance of SAVERCAT in aligning batches in the low-dimensional representation of cells, a task for which many methods exist, is evaluated and benchmarked against existing state-of-the-art software. Next, we compare SAVERCAT to Seurat [6] and scVI [30] in correcting for batch in the high-dimensional gene expression space. We show that SAVERCAT effectively corrects for batch and library size while denoising the data, improving upon existing methods. Finally, we illustrate how SAVERCAT can be applied in complex experimental designs through a single cell sequencing study where directly aligning the batches is not desirable.

## Results

### Model and Methods Overview

Consider the observed scRNA-seq gene expression matrix ***Y*** with *C* rows and *G* columns, where each row represents a cell and each column represents a gene. Here, our model is motivated by the noise properties of UMI-based experiments, but in the next sections we show that SAVERCAT also performs well compared to existing methods on non-UMI-based experiments. Let ***B*** represent the *C* × *l* observed covariate matrix, which can include cell-specific covariates such as library size or sample-specific covariates such as batch, age, sex, etc. We assume ***B*** has been standardized to have column mean 0 and column variance 1. The observed count *Y*_*cg*_ is assumed to follow a Poisson-gamma distribution

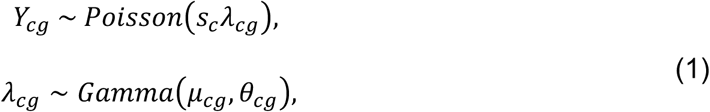

where *λ*_*cg*_ represents the true relative expression that we aim to estimate, *s*_*c*_ is the cell-specific size factor, and *μ*_*cg*_ and *θ*_*cg*_ represent the mean and shape parameters, respectively, of the prior gamma distribution. This Poisson noise model for UMI-based data has been well established [33, 34]. We model the prior mean ***μ***_*c*_ = (*μ*_*cg*_: *g* = 1, …, *G*) as a function *g*_***μ***_ of the cell’s observed covariates ***B***_*c*_ = (***B***_*cj*_: *j* = 1, …, *l*) and its unobserved latent state ***Z***_*c*_, which is a low-dimensional representation of the cell after adjusting for the effect of ***B***_*c*_: ***μ*** = *exp* (*g*_***μ***_ (***Z, B***)).

Our goal is to learn the low dimensional embedding ***Z*** and the function *g*_***μ***_(***Z, B***) which will be used to compute the covariate adjusted and denoised expression matrix. The first step is to construct the low dimensional representation ***Z*** (Fig. 1a). We use a deep neural network, specifically the conditional variational autoencoder (CVAE) [32], to perform nonlinear dimension reduction on the data matrix ***Y***. At this step, we only work with the highly variable genes, see Methods for details. We place a multivariate normal prior distribution on the latent vector *z*_*c*_ for each cell *c* and characterize the variational posterior distribution *q*(*z*_*c*_|***y***_*c*_, ***b***_*c*_) through an encoder function *f*. The variational posterior is thus also a multivariate normal distribution with mean 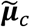 and variance 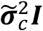. The decoder functions 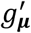 and 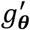 are jointly estimated to characterize the data generating negative binomial distribution *p*(***y***_*c*_|*z*_*c*_, ***b***_*c*_, *s*_*c*_) through the mean 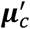 and dispersion 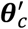. To encourage ***Z*** to capture variation that cannot be captured by ***B***, we train this network by optimizing the weighted objective function

**Figure 1:**
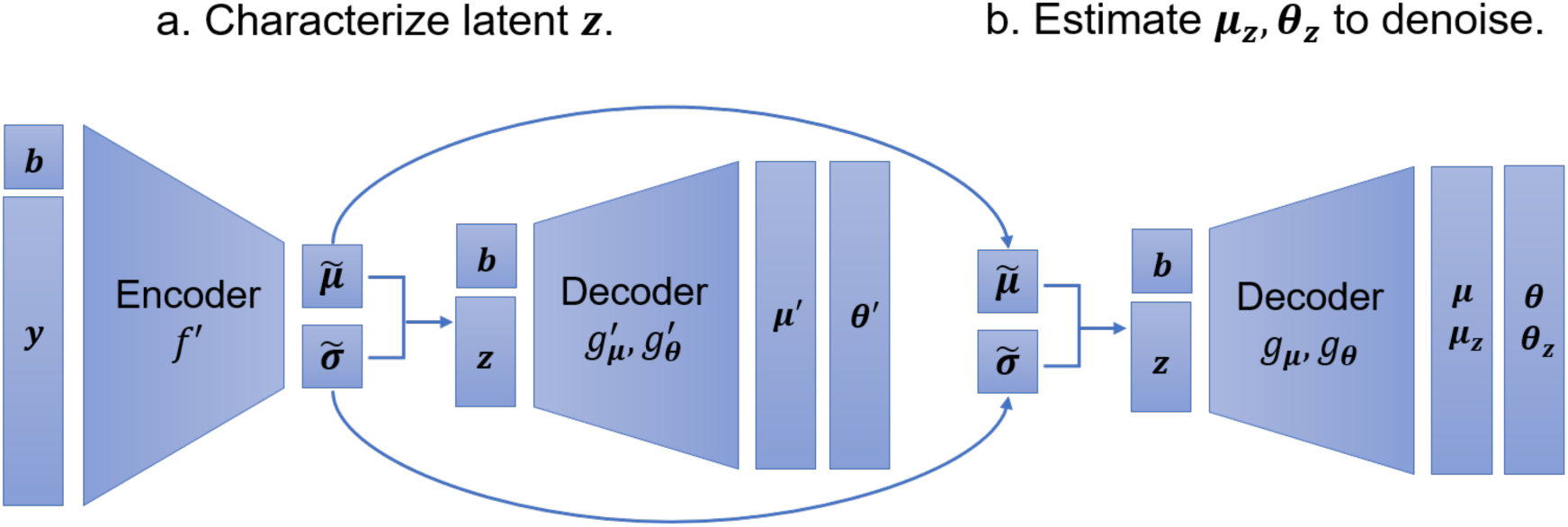
SAVERCAT model. a) In the first step, the conditional variational autoencoder is trained using a subset of highly variable genes and a modified variational loss function to get the latent vector z. b) In the second step, the estimated 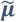 and 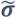 from the first step are used to train the decoder for all genes. **B** can then be set to **0** to estimate the negative binomial parameters **μ**_z_ and **θ**_z_ which characterize the expression of the cells after adjusting for **B**.

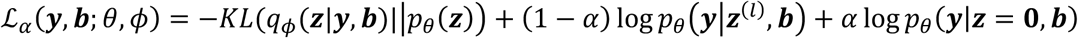

which has a new term *p*(***y***_*c*_|*z*_*c*_ = 0, ***b***_*c*_, *s*_*c*_) in addition to the log-likelihood and Kullback-Leibler divergence terms that are standard in variational autoencoder training. We train the network until the objective function has fully converged (Materials and Methods).

Once the parameters 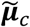 and 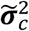 characterizing the cell’s latent representation *z*_*c*_ have been estimated for all cells, we “decode” the latent representation to compute the denoised gene expression matrix (Fig. 1b). For this, we perform a second stage of training, where the output layer is expanded to include all genes. More details are in Methods. The output of the network are the matrices ***μ*** = *g*_***μ***_(***Z, B***) and *θ* = *g*_*θ*_ (***Z, B***) characterizing the prior for the true expression ***λ*** in (1). Now, to remove the effects of *B* on ***λ***, we set ***B*** = 0 as input to the decoder to get ***μ***_*z*_ = *g*_***μ***_ (***Z***, 0) and *θ*_*z*_ = *g*_*θ*_(***Z***, 0). Effectively, this decodes the transcriptome of each cell at the average value of the ***B*** and produces covariate-adjusted parameters for the negative binomial distribution characterizing ***λ***.

The strategy employed by most autoencoder-based denoising schemes [28, 30] is to use the output ***μ*** from the last network layer as the estimate for ***λ***. Similarly, one could use ***μ***_*z*_ as an estimate for covariate adjusted ***λ***, which we denote by ***λ***_*z*_. However, evaluations of denoising methods [24, 35] reveal that directly using the output layer tends to overfit the data even with *k*-fold cross-validation. To combat this, we showed in [24, 27] that, in the case where we don’t adjust for covariates, Bayesian posterior averaging of ***Y*** and ***μ*** substantially improves the estimation of ***λ***. This posterior averaging process can be viewed as a hedging of bets between the model prediction ***μ*** and the observed count ***Y*** – for entries in the matrix where ***μ*** is a poor estimate or where we expect the noise in ***Y*** to be low, our final estimate relies more on ***Y***, and in the opposite scenario, our estimate relies more on ***μ***. To perform Bayesian averaging on our covariate adjusted values ***μ***_*z*_, we need the corresponding count matrix under the setting ***B*** = 0, which is, of course, not observed. Thus, we derive a covariate-adjusted count matrix ***Y***_*z*_ by performing CDF matching of ***Y*** ∼ *NegBinom*(***sμ***, *θ*) and ***Y***_*z*_ ∼ *NegBinom*(***μ***_*z*_, *θ*_*z*_), as follows:

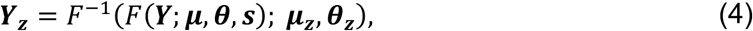

where *F* is the CDF of the negative binomial distribution. The underlying intuition is that we want the distribution of the observed counts ***Y*** given the predicted ***μ*** and *θ* to be identical to the distribution of the adjusted counts ***Y***_*z*_ given the adjusted ***μ***_*z*_ and *θ*_*z*_, preserving the technical noise characteristics of scRNA-seq expression count data. Then, the empirical Bayes SAVERCAT posterior for ***λ***_*z*_ given ***Y***_*z*_ is

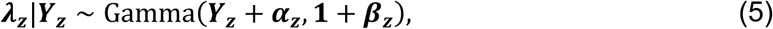

where ***α***_*z*_ and ***β***_*z*_ are the shape and rate parameters of the prior gamma distribution reparameterized from ***μ***_*z*_ and *θ*_*z*_. The mean of this posterior distribution is our SAVERCAT covariate-adjusted and denoised estimate of gene expression. Sampling from this posterior distribution allows quantification of the estimation uncertainty.

Hence, the SAVERCAT software package outputs three values for each gene in each cell: (1) The covariate-adjusted counts ***Y***_*z*_, which mimic what you would have observed if you performed the experiment in the setting ***B*** = 0 (e.g. all cells in same batch) and sequenced at the same coverage, (2) the denoised estimate 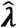, which mimics what you would have observed if you performed the experiment in the setting ***B*** = 0 but with higher sequencing coverage, and finally, (3) a sample from the posterior distribution (equation 5). These three outputs are useful for different types of downstream analyses. Based on evaluations in [36], we recommend performing clustering, trajectory, and visualization analyses using the denoised estimates, differential expression testing using the covariate-adjusted counts, and characterization of gene expression distributions using the Bayesian posterior sampled values.

### Evaluation of batch correction in low dimensions

We start by evaluating performance on the problem of constructing a low-dimensional representation of the cells correcting for batch, for which many methods have been developed. Recently, Tran et al. [37] performed an in-depth comparison of 14 batch correction methods, in which they found Seurat [6], Harmony [8] to have good overall performance. We thus adopt the benchmarking data sets and metrics in Tran et al. [37] and compare our method with Seurat [6], Harmony [8], and scVI [30]. scVI was not assessed in [37], but we include it here because it is also a newer method based on deep neural networks. Implementation details for each method are provided in Materials and Methods. The benchmarking datasets from [37] cover a diverse collection of cell types such as immune cells, pancreas cells, retina cells and brain cells and all of the main technologies such as Smart-Seq, 10X, MARS-Seq and Drop-Seq (Materials and Methods). Both quantitative evaluation with local inverse Simpson’s index (LISI, Materials and Methods) [8] and visualization using UMAP and t-SNE (Supplementary Note 3) were applied to assess method performance on each data set. In evaluating each method, it is important to note the inherent trade-off between cell type separation (1-cLISI) and batch mixing (iLISI), in the sense that aggressive batch correction leads to a loss of cell types signatures, when these signatures are not independent of batch. As a result, methods that are more aggressive in mixing batches, with higher iLISI, may reduce the separation between cell types and thus have a lower 1-cLISI. For example, in dataset 1, SAVERCAT was the top method in terms of batch mixing, but the worst in separating cell types (Fig. 2).

**Figure 2:**
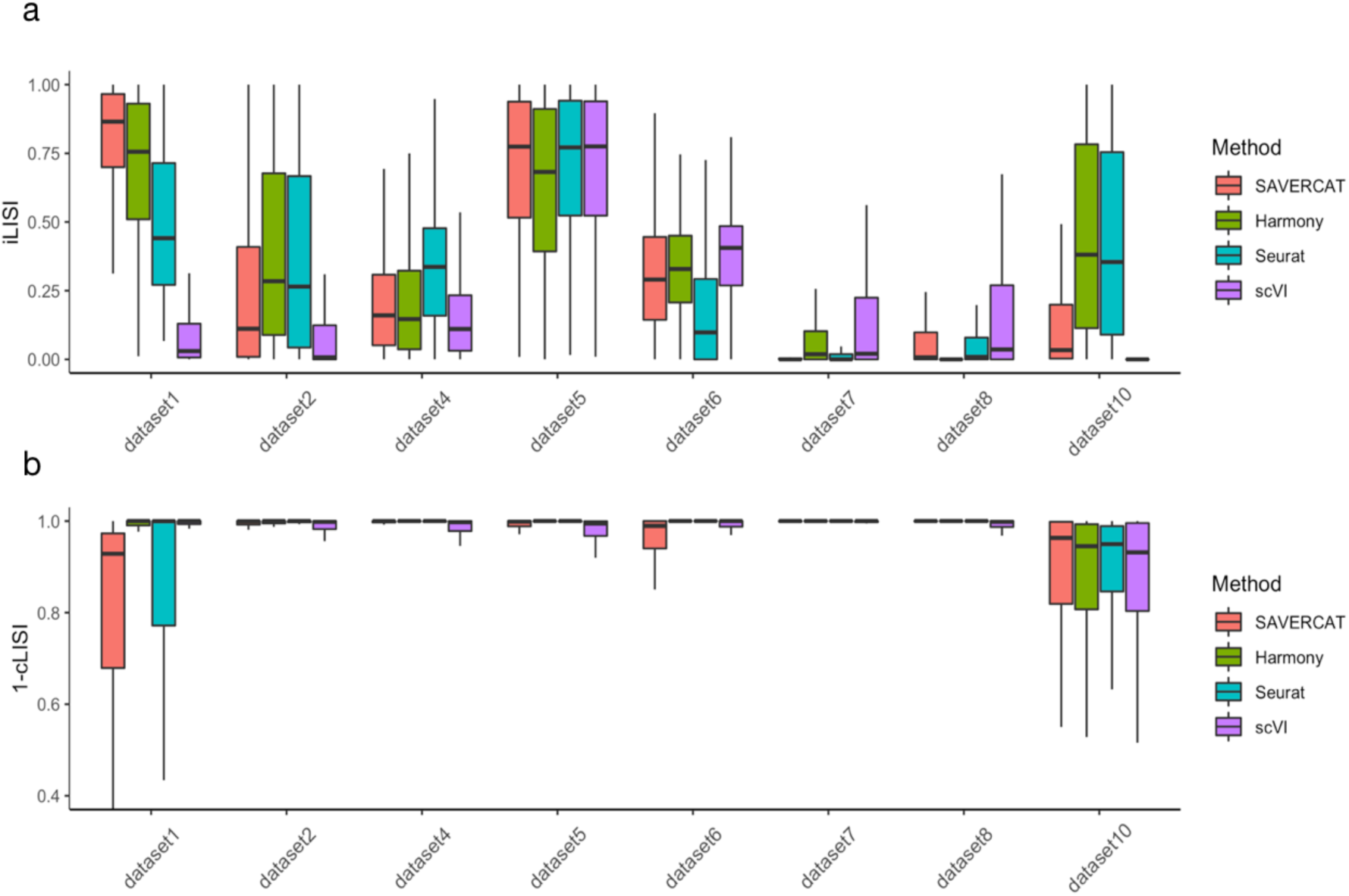
Comparison of SAVERCAT with Seurat, Harmony and scVI in removing batch effects in the low-dimensional latent representation using LISI scores. a) A higher iLISI value reveals more uniform batch mixing. b) A higher 1-cLISI value reveals better separation between cell types. Both iLISI and cLISI are calculated with `lisi` package and then scaled to 0 and 1.

We found that all four methods performed well across datasets in terms of cell type separation measured by 1-cLISI (Fig. 2). There are clear differences between data sets in terms of the inherent difficulty of batch correction (Fig. 2), and no method wins every time. SAVERCAT performed well compared to the best of the field, even in scenarios where the number of cells is small for training a neural network (dataset 1). For datasets in which there are cell types unique to specific batches (dataset 1, 4, 7, 8 and 10), our method is capable of preserving the distinct cell types when removing the batch effect.

### Correcting for batch in the gene expression matrix

In the previous section, we showed that SAVERCAT performs comparably to the best of existing methods in removing batch effects, while maintaining biological signals, in the *low-dimensional representation* of the cells. Now we consider the problem of removing batch effects in the original gene expression matrix, for which fewer methods exist and no rigorous third-party benchmarking is available. First, consider the 10x PBMC datasets from three healthy donors sequenced using three different 10x technologies – VDJ, V2, and V3 {n.d., Datasets - 10x Genomics}. Dataset 5 in the previous section used the V2 and V3 PBMC datasets. We chose this data set because it is the most successful case-study for batch-correction in low dimensional representation, where all four methods (SAVERCAT, Harmony, Seurat, and scVI) were able to obtain a good mixing between batch while maintaining the distinction between cell types. Only with good batch mixing in the low-dimensional representation can we meaningfully explore the effectiveness of batch correction in the original gene expression matrix. For the PBMC dataset, we identified three main cell types – monocytes, T/NK cells, and B cells – which separated well according to the UMAP visualization (Supplementary Fig. 1). Each technology was treated as a batch and our goal is to align the gene expression values across batches *within* each main cell type. Since Harmony does not output a batch-corrected gene expression matrix, we focused on SAVERCAT, Seurat, and scVI in this analysis. For SAVERCAT, we considered both the batch-corrected count output and the posterior sampled output.

First, we took the batch-corrected gene expression matrices from each method and performed dimension reduction and UMAP visualization on a set of highly variable genes following the Seurat clustering workflow (Fig. 3a). In the original dataset, the cells separated according to both batch and cell type, resulting in nine distinct clusters in the UMAP visualization. Seurat, scVI, SAVERCAT counts, and SAVERCAT denoised output all produced cell clusters which are separated by cell type and well-mixed by batch.

**Figure 3:**
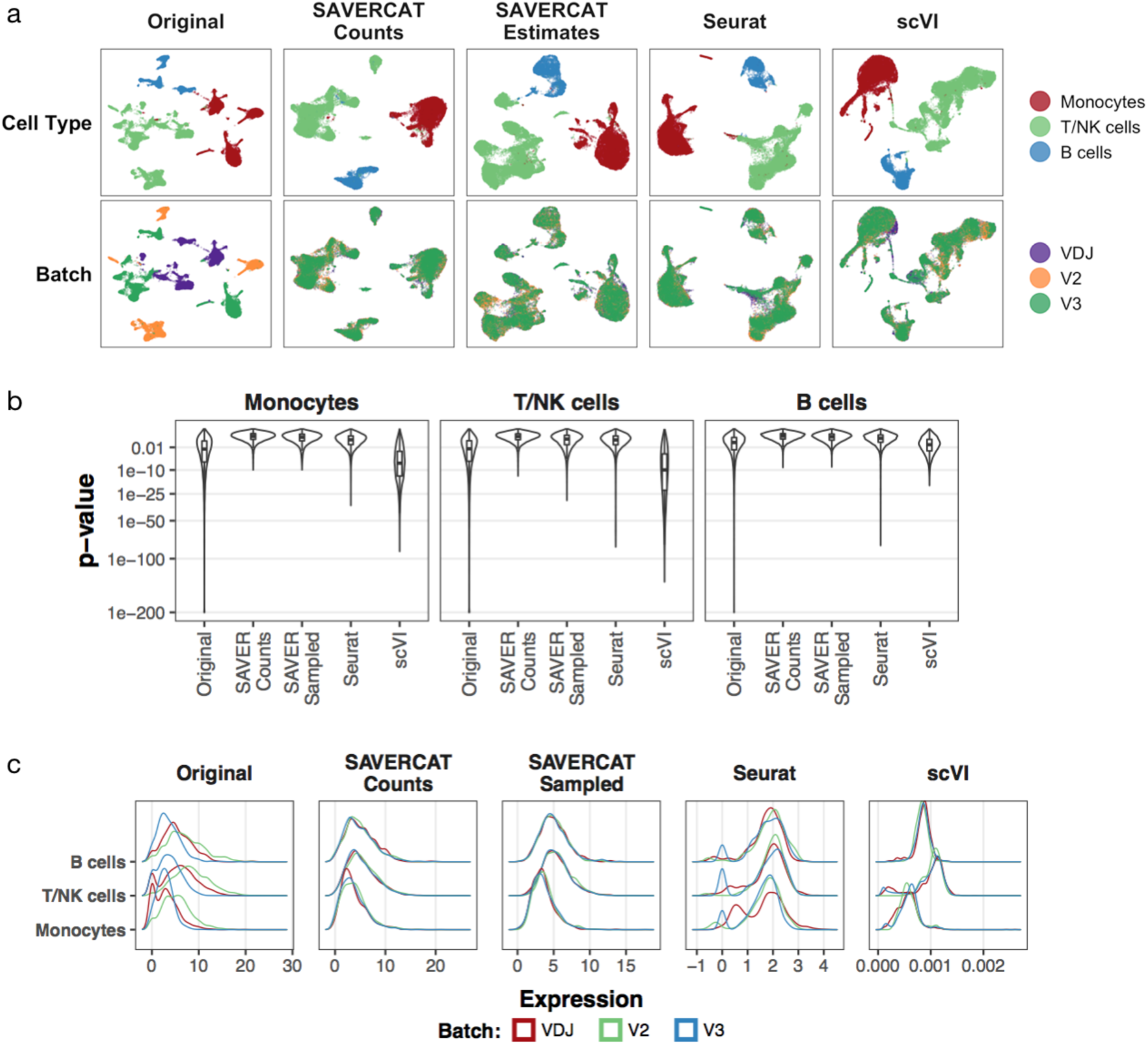
Correcting for batch in the gene expression matrix in the PBMC dataset. a) UMAP visualization of original and batch-corrected expression matrices. b) Violin plots of p-values from Kruskal-Wallis tests for differences in distribution between batches within each main cell type for each gene. c) Kernel density plots of gene expression for gene HNRNPA1.

Next, we focused on the batch corrected expression matrices and conducted differential expression test for each gene, testing for difference between batches within each cell type and across all cells. We considered three statistical tests: one-way ANOVA, the Kruskal-Wallis test [38], and the Anderson-Darling *k*-sample test [39]. The one-way ANOVA is a parametric test for differences in mean expression between groups, while the Kruskal-Wallis test and the Anderson-Darling *k*-sample test are non-parametric tests for differences in distribution of expression between groups. Applying the one-way ANOVA to the original data revealed that while the majority of genes have similar distributions across batches within each cell type, there are a few genes which exhibit strong batch effects, as indicated by a low *p*-value (Fig. 3b). 16.7% of genes have a *p*-value less than 10^−10^ for the monocyte population, 16.0% for the T/NK cell population, and 5.5% for the B cells (Supplementary Table 1). The SAVERCAT counts and sampled posterior values significantly reduce the effects of batch with the vast majority of genes having *p*-value greater than 0.01. Seurat also performs well although with a larger proportion of genes than SAVERCAT testing highly significant. On the other hand, the median *p*-value for scVI is lower than that of the original data for all cell types, indicating that scVI does not effectively remove batch effects for most genes.

The *k*-sample Anderson-Darling test and the Kruskal-Wallis test applied to the original data revealed the same pattern of a small proportion of genes driving the batch effect (Supplementary Fig. 2, Supplementary Table 1). To examine the batch-corrected expression distributions, we also plotted the kernel density curves of expression estimated for each batch within each cell type (Fig. 3c, Supplementary Fig. 4) for a selection of cell type markers. Accurate batch correction would result in identical expression distributions across the three batches, while maintaining the true expression differences between cell types. These results show that SAVERCAT effectively removes the difference between batches at the gene level. In contrast to its performance as evaluated by one-way ANOVA, Seurat performs quite poorly in aligning gene expression distributions, with more than 96% of genes testing highly significant for the *k*-sample Anderson-Darling test. Seurat appears to effectively remove the mean batch effect but introduces differences in distribution, as evident by its low p-values in the non-parametric distribution tests, and alsovisible in the density curves in Figure 3c and Supplementary Fig. 4.

**Figure 4:**
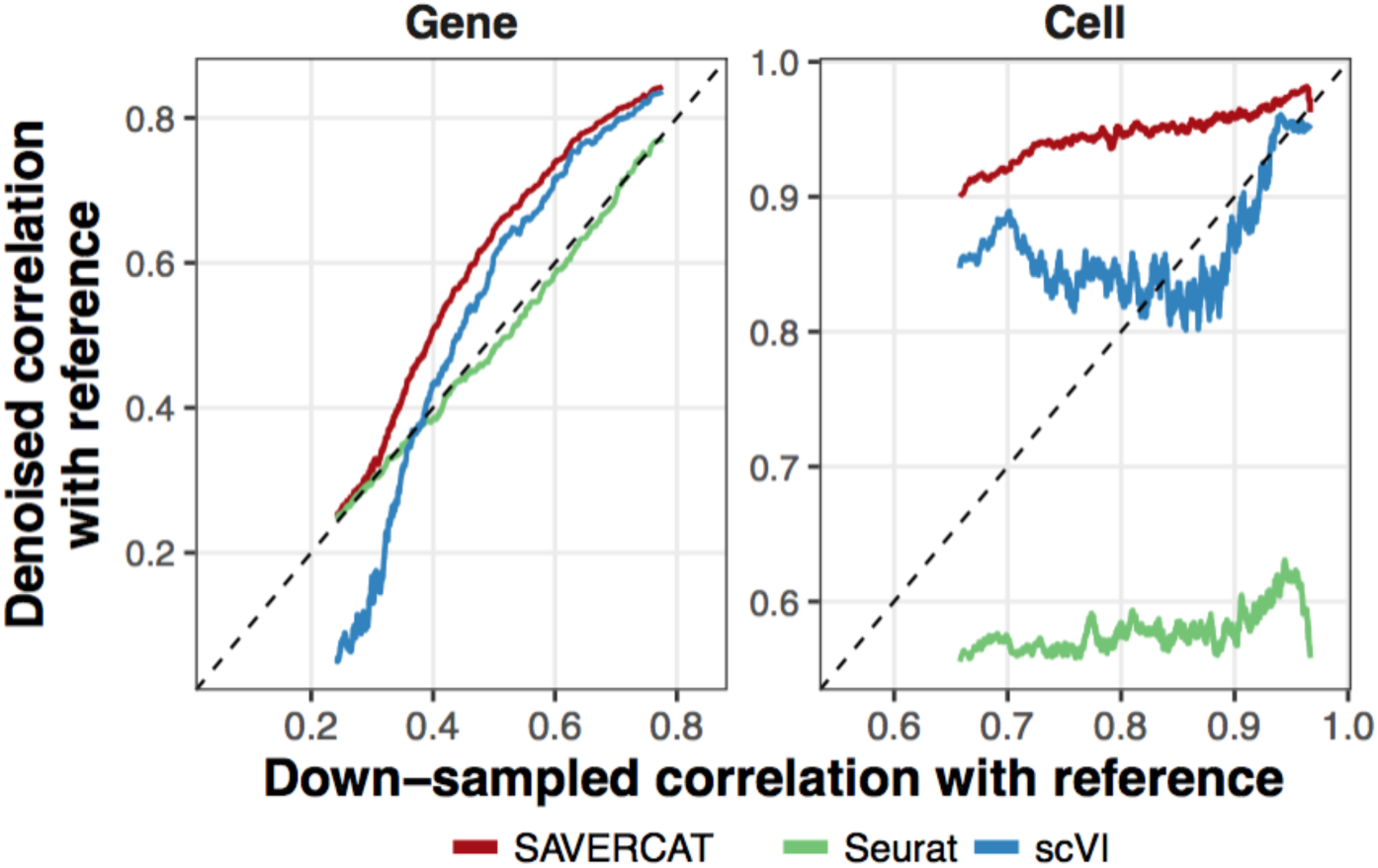
Comparison of down-sampled correlation and denoised correlation with reference across genes and cells.

We also calculated the correlation between the corrected expression of each gene with the log_10_ library size (Supplementary Fig. 3). Since library size is a technical covariate, low absolute correlation with expression is desired. SAVERCAT covariate-adjusted counts and SAVERCAT denoised values all have correlations with library size that is centered at zero, and more densely concentrated around zero than what is achieved by a simple library-size normalization. The Seurat-corrected expression exhibits slightly positive correlation with library size while scVI expression is highly positively correlated with library size for some genes while highly negatively correlated with library size for others.

### Integration of batch correction and denoising in gene expression recovery

The previous sections assess the effectiveness of methods in removing the influence of observed confounding covariates on gene expression. Now, we turn to denoising, which removes technical variation that is due to the inherent random sampling introduced in the library preparation and sequencing steps of the experiment [33, 34]. Since effective denoising should mimic what we would get if we sequenced the cDNA libraries at higher depth, and thus give gene expression estimates that are closer to the truth, denoising accuracy is often evaluated through down-sampling: The reads of a scRNA-seq dataset are randomly sampled to a lower coverage, the denoising/preprocessing method is applied to the lower coverage data set, and the denoising output is then compared against the original data matrix. Since SAVERCAT performs denoising simultaneously with batch correction, we extended this down-sampling procedure to evaluate the effectiveness of batch-corrected denoising: Consider the 10x PBMC dataset as an example, we selected 3,534 genes and 4,461 cells from the V3 batch to serve as our reference data with no batch effects. We then randomly assigned each cell to a batch (VDJ, V2, or V3) and added batch effects by perturbing the observed value for each gene in each cell cell by the average cell-type specific batch effect in the original 10x VDJ, V2, and V3 datasets (Materials and Methods). We then performed down-sampling on this batch-perturbed matrix to get the “observed” expression matrix with batch effects. The effectiveness of batch-corrected denoising is evaluated by calculating the gene-wise and cell-wise correlations of the batch-corrected denoised dataset with the reference dataset.

Results are shown in Figure 4, where gene- and cell-wise correlations are plotted against the original gene- and cell-wise correlations with reference of the down-sampled data set prior to batch-correction and denoising. The original correlations are a measure of the degree of technical noise in the gene. By gene-wise correlation, SAVERCAT improves correlation with reference for all genes. This is due to the empirical Bayes shrinkage and cross-validation steps in SAVERCAT which enables it to be at least as accurate as the original counts for each gene, and not “sacrifice” some genes for the sake of global performance measures. Even though Seurat is not a denoising method, we included it here because it is a popular pipeline. We see that its gene-wise correlation with the reference is similar to that of the original down-sampled dataset. scVI improves the gene-wise correlation for genes which have moderate or high correlations with the reference to a lesser extent than SAVERCAT but decreases the correlation for the noisy genes that have low correlation with the reference. We believe this is due to overfitting from using the predicted values directly without shrinkage. In terms of cell-wise correlation, SAVERCAT substantially improves the correlation with the reference for all cells while Seurat, surprisingly, decreases the correlation with reference. This can be explained by Seurat’s corrected values not being on the same scale as the original expression for each gene, which results in lower correlations. scVI increases the correlation with the reference for cells with low observed correlations with the reference but performs similarly as cells with moderate or high correlations with the reference. We also assessed the effectiveness of batch correction, via the same metrics as in the previous section, on this down-sampled dataset and discovered the same trends as in the original PBMC dataset (Supplementary Fig. 5).

**Figure 5:**
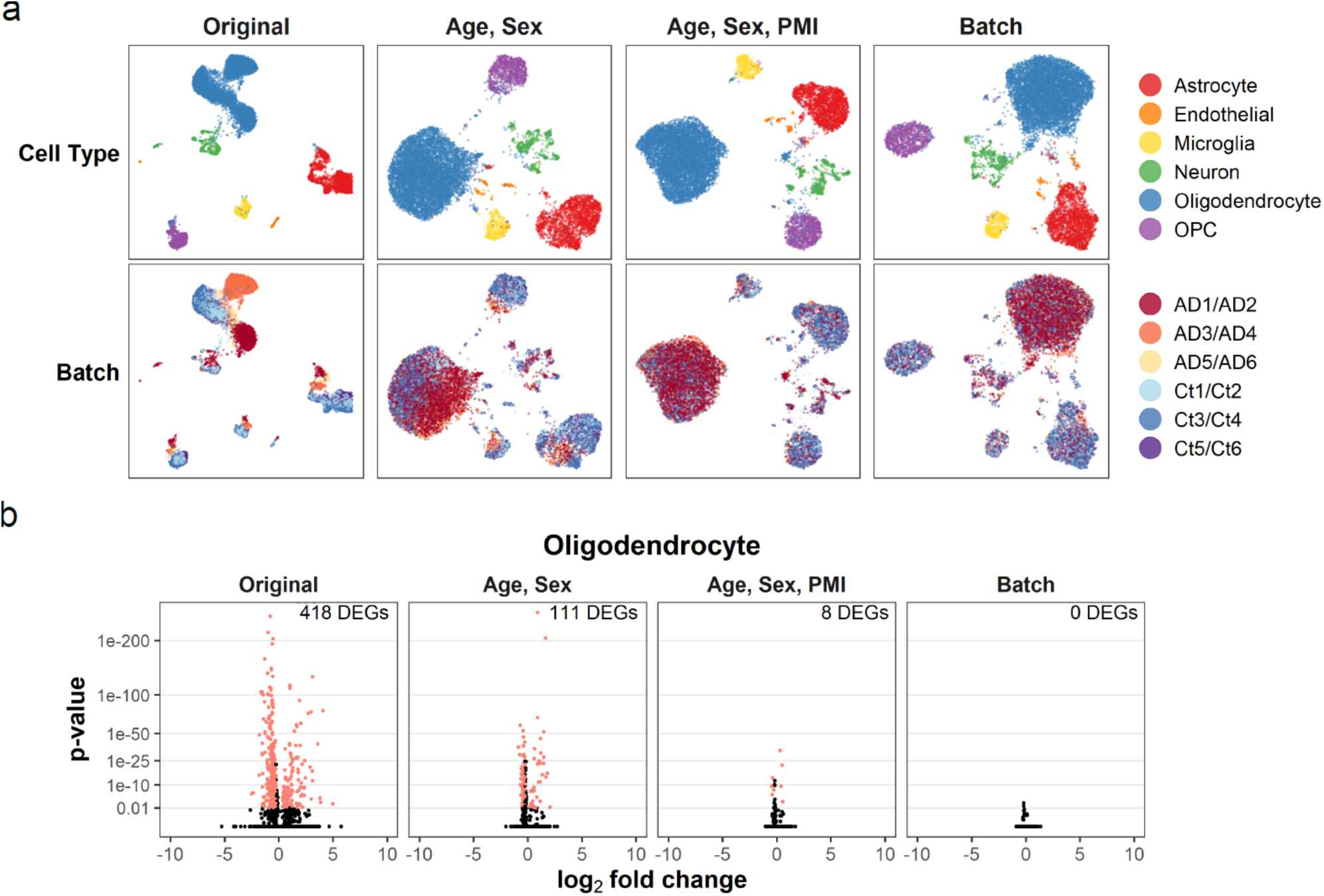
SAVERCAT applied to Alzheimer’s dataset. a) UMAP visualization of original data and SAVERCAT low-dimensional representation under different adjustment scenarios. b) Volcano plots of adjusted Wilcoxon rank-sum test p-values of gene expression and log_2_ fold change between Alzheimer’s cells and control cells with number of DEGs discovered for the original data and SAVERCAT counts under different adjustment scenarios.

### Alzheimer’s case study

The analyses in the preceding sections have shown that, given a set of batch labels, SAVERCAT can learn a low-dimensional latent representation of the data and denoise gene expression while correcting for batch. However, in real applications, directly aligning all of the batches may not be desired if batch is perfectly confounded with a variable of interest. In some studies there are also additional covariates, such as subject age or sex, which we hope to account for. We now apply SAVERCAT to one such study on Alzheimer’s disease from Grubman et al. [13] to demonstrate how it can flexibly accommodate a more complex experimental design and adjust for both continuous and discrete covariates. In Grubman et al. [13], single nucleus RNA-seq data was collected from the entorhinal cortex region of the brains of six human patients with Alzheimer’s disease and six control subjects. Patient characteristics can be found in Supplementary Table 3.

Grubman et al. studied the differences in cell-type specific gene expression between Alzheimer’s patients and control subjects and identified differentially expressed genes (DEGs). However, there are confounding covariates which may affect the analysis. For example, although an attempt was made to match the Alzheimer’s patients and controls by age and sex, perfect matching is impossible and there are still discrepancies in these two variables between the two groups. Furthermore, patients with Alzheimer’s disease have substantially shorter post-mortem interval (PMI) than the control subjects. The most glaring possible confounder is batch. For each batch, two patients were sequenced but they were either both Alzheimer’s patients or both control. Thus, batch is completely confounded with the question of interest --- the difference between Alzheimer’s patients and control subjects. This type of confounding is often unavoidable in single cell studies involving human subjects, where the access to samples and timing of experiments is hard to control. Due to this perfect confounding, naively aligning samples across all batches is not desirable, as the goal of batch correction methods is to remove, through a continuous transform, local differences between batches that underlie cell-type specific DEGs. Thus, a more careful consideration of which variables to adjust for and how it would affect the interpretability of downstream results is necessary.

We applied SAVERCAT to this data under three adjustment schemes, ranging in severity of adjustment. In the most conservative adjustment scenario, we adjusted for age and sex. Since age and sex are less correlated with diagnosis, as compared to PMI and batch, we expect adjustment by these variables to preserve more of the meaningful biological signal. In the moderate adjustment scenario, we adjusted for PMI in addition to age and sex. The risk of adjusting for PMI in this experiment is that it is strongly correlated with diagnosis, and thus we may erase meaningful biological signal. Finally, in the most aggressive adjustment scenario, we adjusted directly for batch, which is quite standard in current single cell analysis pipelines. In all scenarios, we also adjusted by library size. Grubman et al. did not attempt to remove the effects of batch or any of these potentially confounding variables in their analysis of the data, most likely due to the confounding between batch and diagnosis.

First, we visualized the latent representation of the cells using the original expression matrix and the covariate-adjusted SAVERCAT latent matrix under the three scenarios described above. For each case, the cells are colored in two ways: by cell type and by batch (Fig. 5a). The cell type labels were identified by Grubman et al. using cell type-specific marker genes. For the original data, cells generally cluster according to cell type. However, within cell types, especially oligodendrocytes and astrocytes, there are well-separated clusters of cells which are explainable by batch, even when limited to cells from the control patients. Thus, it is hard to disentangle batch effect from the separation between controls and Alzheimer’s patients within each cell type. When we look at the visualization of SAVERCAT after adjustment of age and sex, we see that while the cell types still separate cleanly, there is now a clear mixing of the batches by diagnosis within each cell type. For example, the cells in batch AD1/AD2 colored red and the cells in batch AD3/AD4 colored orange are completely separated in the original data for oligodendrocytes. After adjusting for age and sex, the cells in these two batches are visually very well mixed. Thus, adjusting for age and sex allows us to align the batches within each cell type without explicitly doing batch alignment, while still retaining some of the differences between disease and control.

Now consider the SAVERCAT result where, in addition to age and sex, we also adjust for PMI, we see that the distinction between cell types is preserved, but all six batches are aligned. There is no more separation between Alzheimer’s disease patients and control subjects. Thus, adjusting for PMI, which is highly correlated with diagnosis, removes a substantial amount of variation which was previously attributed to differences in diagnosis. Finally, when we explicitly correct for batch cells from each batch and diagnosis become well mixed and there is almost no difference between disease and control cells. We also evaluated the prevalence of batch effects in the original and SAVERCAT-corrected gene expression matrix and discovered that as the severity of the adjustment increases, gene expression distributions between batches within each disease condition become more similar (Supplementary Fig. 7, Supplementary Table 4).

We conducted differential expression analysis between cells collected from Alzheimer’s patients and cells collected from control subjects within each cell type. We applied the non-parametric Wilcoxon rank-sum test for the original expression counts and the three SAVERCAT adjusted count matrices. Taking the oligodendrocytes as an example, we discovered 418 DEGs using the original expression counts (Fig. 5b). However, due to the clear batch effects in the oligodendrocyte population even among cells from the control group, it is unclear how many of the 418 DEGs discovered are due to differences in batch. Differential expression analysis on the age and sex adjusted SAVERCAT counts yielded 111 DEGs at a lower overall significance and lower fold change difference than the original expression counts. Adding PMI as a covariate substantially reduced the number of DEGs from 111 to only 8. Since PMI is highly correlated with diagnosis in this experiment, adjusting for PMI effectively removes most of the differences in gene expression between Alzheimer’s and control cells. Unsurprisingly, adjusting directly for batch essentially removed all differences between patients and control. The same trend was observed for the other cell types (Supplementary Fig. 8).

In summary, explicitly correcting for batch eliminates all of the relevant biological differences in this study, while not doing any correction results in a substantial number of differentially expressed genes which may be false positives. With SAVERCAT, we explored two intermediate adjustment schemes, and found that correcting for age and sex effectively aligns the batches among the control samples, and still retains differences between disease and control cells.

## Discussion

With the broad adoption of single cell sequencing in population-wide and clinical studies, removing the effects of confounding subject- and cell-level covariates has become a pervasive problem. While many methods have been developed for batch alignment, these methods are difficult to apply when batch is confounded with the variable of interest, or when there are other confounding variables within each batch. A further difficulty when applying existing batch alignment methods is that while they can be effective in removing batch effects in the low-dimensional representation of cells, it is much more difficult to remove batch effects, and adjust for covariates, in the original high-dimensional gene expression matrix. In this work, we presented SAVERCAT, a framework for covariate-adjusted representation learning and denoising of single-cell RNA-seq. SAVERCAT is comparable to the top-performing batch correction methods in removing the batch effect in the low-dimensional latent representation, and improves upon existing methods in removing batch effects and other confounding covariates, such as cell library size, from the expression matrix. In addition to covariate adjustment, SAVERCAT also denoises the data, thus reducing the technical noise due to inherent random sampling. In addition to the denoised and batch-corrected expression matrix, SAVERCAT also gives the posterior distribution of each entry in the expression matrix, which reflects the uncertainty of the estimate for each gene in each cell.

Directly aligning all batches is oftentimes not the best approach especially in clinical and experimental settings where batch cannot be randomized with respect to the contrast of interest. A more careful and systematic consideration of how observed technical covariates affect an analysis needs to be conducted. The choice of which variables to correct for depends largely on the context of the experiment. In the Alzheimer’s data, we saw how correcting for post-mortem interval, a variable which is correlated with diagnosis, improves the mixing of batches but removes almost all differentially expression genes. Correcting for age and sex, on the other hand, allows for the mixing of batches within diagnosis group and still preserves some differential expression signals. In situations like this, it will need to be up to the researcher to determine how stringent the correction should be. Although we have shown the capability of our method to adjust for observed covariates, a balanced study would ultimately lead to more interpretable and reproducible findings.

## Materials and Methods

### CVAE with weighted objective

SAVERCAT is based on the conditional autoencoder, which uses a neural network to capture the generative process of the data ***y*** from latent variable *z* and observed variable ***b*** [32]. We introduce a modified objective function to encourage *z* to capture separate sources of variation not captured through ***b***. In the variational autoencoder framework, the latent variable *z* is generated from a prior distribution *p*_*θ*_ (*z*) and the output ***y*** is generated from the distribution *p*_*θ*_ (***y***|*z*, ***b***). The true posterior distribution *p*_*θ*_(*z*|***y, b***) is usually intractable so a parametric distribution *q*_*ϕ*_(*z*|***y, b***) is used to approximate the posterior. In the CVAE, *q*_*ϕ*_(*z*|***x, b***) is the encoder and *p*_*ϕ*_(***y***|*z*, ***b***) is the decoder. The standard empirical objective of the variational lower bound is

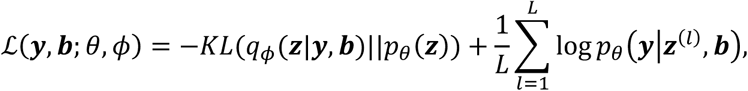

where *z*^(*l*)^ = *f*_*ϕ*_(***y, b***, *ϵ*), *ϵ* ∼ *p*(*ϵ*) and *f*_*ϕ*_(·) is a deterministic encoder function used in the reparameterization trick. For the CVAE used in SAVERCAT, we let *L* = 1 and *ϵ* ∼ *𝒩*(0, *I*). To encourage *z* to capture separate sources of variation not captured through ***b***, we define the weighted empirical objective as

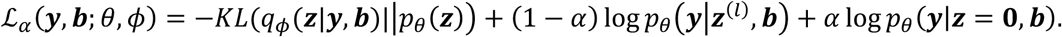

Placing a positive weight on *log p*_*θ*_ (***y***|*z* = 0, ***b***) encourages the network to reconstruct ***y*** using only information contained in ***b***. This allows the characterization of *z* as latent variables which capture variation in ***y*** not explained by ***b***.

### SAVERCAT implementation details

The SAVERCAT method consists of two main steps: (i) Finding a low-dimensional latent representation of the cells while adjusting for observed covariates and (ii) finding a covariate-adjusted count matrix and denoised output. Let *s*_*c*_ be the size factor defined by 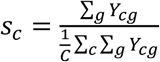. In the first step, we train the CVAE using the standardized log expression 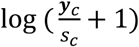 and observed covariate vector ***b***_c_ as input to the encoder function *f′*. By default, we calculate batch-level highly variable genes using Seurat’s FindVariableFeatures function and select the top 3000 genes from the union across batches. The output of the encoder function *f′* is the multivariate normal distribution mean parameter 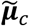 and the variance parameter 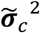. By default, the dimension of the multivariate normal is set to be 30. *z*_*c*_ is sampled from the multivariate normal distribution and is concatenated with ***b***_*c*_ as input to the decoder function 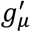. to get the negative binomial mean 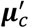. To model the negative binomial dispersion 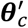, we feed ***b*** _*c*_ as input to the dispersion decoder function 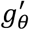. Here, we assume that the dispersion is a function of only the external covariates. The output of the decoder is the vector of negative binomial means 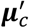 and dispersions 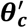, where the data generating distribution is 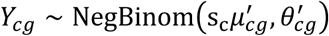. Then the weighted empirical objective function is

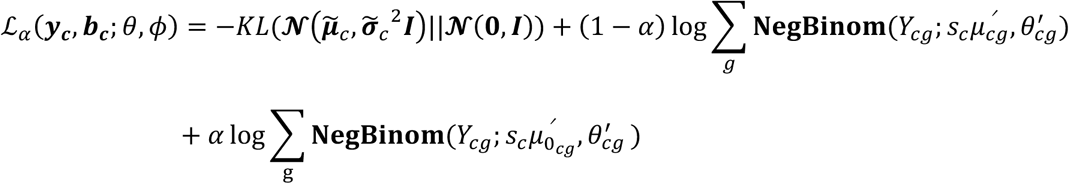

where 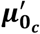 is the output of the decoder 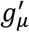 concatenated with *z*_0_ = 0 as the input. The objective function is optimized using the Adam optimizer [40], setting *α* = 0.01. The encoder and decoder networks both consist of two fully-connected 128-node layers with the LeakyReLU activation function. Batch normalization [41] without the learnable shift parameter is applied after the activation functions of the encoder layers. It is important to remove the shift parameter to ensure that the output of the layers have mean 0 since the weighted objective function relies on *z* being centered. Dropout [42] is used in the first encoder layer with a dropout fraction of 0.1. KL annealing is performed during the start of training and training is stopped once the objective function has converged. The learned multivariate normal mean vector 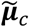 is a low-dimensional representation of cell *c* and can be used in visualization and clustering analyses.

The second step of SAVERCAT is denoising. We take the estimated 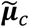and 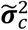 and train another decoder network with functions *g*_***μ***_ and *g*_*θ*_. Like before, we sample *z*_*c*_ from the multivariate normal distribution characterized by 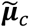 and 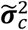 and concatenate with ***b***_*c*_ as input to the decoder network. In matrix notation, the output is ***μ*** = *g*_***μ***_(***Z, B***) and *θ* = *g*_*θ*_(***Z, B***), where the data-generating distribution is *Y*_*cg*_ ∼ *NegBinom*(*s*_*c*_*μ*_*cg*_, *θ*_*cg*_). By default, we train the decoder on all genes. The weighted objective function takes the same form as in the first step but there is no longer a KL-divergence term. *α* is set to 0.01 and the Adam optimizer is used. To prevent overfitting, we perform *k*-fold cross-validation to select the epoch which gives the lowest validation error. We then train using all cells and perform early stopping at that epoch. After training, we generate the estimates of ***μ*** and *θ* by using the multivariate normal means 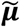 and ***B*** as input to the decoder. To obtain the covariate adjusted ***μ***_*z*_ and *θ*_*z*_, we input ***B*** = 0 to the decoder.

### Benchmarking analysis

The benchmarking datasets, batch labels, and cell type labels were obtained from Tran et al. [37]. Here is a brief description of the datasets:

- Dataset 1: 576 non-UMI human dendritic cells from 2 batches [43]
- Dataset 2: 6,954 non-UMI cells from two Mouse Cell Atlas studies [44, 45]
- Dataset 4: 14,767 non-UMI and UMI human pancreas cells from five studies [46-50]
- Dataset 5: 15,476 UMI human PBMC cells from two 10x Genomics technologies {n.d., Datasets - 10x Genomics}
- Dataset 6: 9,530 UMI human 293T and mouse 3T3 cells in three batches [51]
- Dataset 7: 71,638 UMI mouse retina cells from two studies [52, 53]
- Dataset 8: 100,000 sub-sampled UMI mouse brain datasets from two studies [54, 55]
- Dataset 9: 100,000 sub-sampled UMI bone marrow and cord blood-derived cells from the Human Cell Atlas [56]
- Dataset 10: 4,649 non-UMI and UMI mouse hematopoietic stem and progenitor cells from two studies [57, 58]

SAVERCAT, scVI v0.6.4 [30], Seurat v3.1.5 [6], and Harmony v1.0 [8] were applied to each dataset. SAVERCAT was run with default options for datasets. scVI was run using a subset of 3,000 highly variable genes. Seurat was run following the standard data integration workflow downloaded from github repository (JinmiaoChenLab/Batch-effect-removal-benchmarking) [37] except reciprocal PCA was used for datasets 7, 8, and 9 due to memory issues. Harmony was run following the integration pipelines downloaded from github repository (JinmiaoChenLab/Batch-effect-removal-benchmarking) [37] which sets maximum number of rounds to run clustering to be 100 and number of clusters to be 50 and uses Seurat v2.3.4 for highly variable gene selection. The local inverse Simpson’s index (LISI) [8] was calculated using the *lisi R* package version 1.0 with default *perplexity* = 30. With low-dimensional embeddings calculated from each method, we compute cell-specific LISI scores with respect to batch labels (iLISI) and cell type labels (cLISI) for each dataset separately. iLISI defines the effective number of batches in the neighborhood of a cell which quantifies local batch mixing while cLISI defines the effective number of cell types which quantifies local cell type mixing. For a dataset which has equal number of cells in each batch, iLISI ranges from 1 to the number of batches. For a dataset which has equal number of cells in each cell type, cLISI ranges from 1 to the number of cell types. We scaled the iLISI and cLISI scores for each dataset using the theoretical lower and upper bounds for a balanced dataset so that it ranges from 0 to 1.

### PBMC original gene expression analysis

The VDJ 5’ dataset of 8,258 PBMC cells was downloaded from https://support.10xgenomics.com/single-cell-vdj/datasets/3.0.0/vdj_v1_hs_pbmc2_5gex_protein. The V2 dataset of 8,381 PBMC cells was downloaded from https://support.10xgenomics.com/single-cell-gene-expression/datasets/2.1.0/pbmc8k. The V3 dataset of 11,769 cells was downloaded from https://support.10xgenomics.com/single-cell-gene-expression/datasets/3.0.0/pbmc_10k_v3. Seurat v3.1.4 [4, 6] was used to cluster the cells within each dataset individually and cell types were identified by monocyte, T/NK cell, and B cell marker genes. Unidentified cells were removed and after filtering out genes with non-zero expression in less than 2 cells, the combined dataset consisted of 22,341 genes and 28,022 cells.

SAVERCAT, Seurat, and scVI were applied to the PBMC original dataset to correct for batch in the expression matrix. SAVERCAT was run with default settings. Seurat was run according to the standard workflow except setting the *features*.*to*.*integrate* parameter in the *IntegrateData* function to all genes. scVI was run using all genes and trained for 200 epochs. The batch-corrected output was obtained by averaging the normalized negative binomial mean across the three batches. The batch-corrected expression matrices were then supplied to the Seurat clustering workflow to perform UMAP visualization. The Anderson-Darling *k*-sample test was performed using the *kSamples* R package version 1.2-9.

### PBMC down-sampling experiment

To create the reference dataset with no batch effects, we used the V3 dataset and selected cells in the top 40^th^ percentile of library size from each of the three cell types and genes with non-zero expression in more than 20% of cells. This resulted in a reference dataset ***λ*** of 3,534 genes and 4,461 cells. To simulate cell type-specific batch effects, we randomly assigned cells in the reference dataset to one of VDJ, V2, or V3, and perturbed the expression by the average cell-specific batch effect in the original PBMC dataset. For example, consider a gene *g* and let 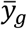 represent the average expression across all cells for gene *g*. Let 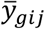 represent the average expression of gene *g* across cells belonging to cell type *i* and batch *j*. We calculate the average cell type-specific batch effect of cell type *i* and batch *j* as 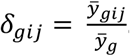. Consider the reference expression *λ*_*cg*_. Then the batch corrected reference expression is 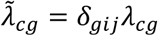 for gene *g* and cell *c* belonging to cell type *i* and batch *j*. We then sample the observed expression as 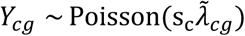, where *s*_*c*_ ∼ *Gamma*(10, 100). We applied SAVERCAT, Seurat, and scVI to the down-sampled ***Y*** and calculated the gene-wise and cell-wise correlations with respect to the reference ***λ***. For the four down-sampled datasets, we performed the down-sampling as described in Huang et al. [24].

### Alzheimer’s analysis

The Alzheimer’s dataset from Grubman et al. [13] was downloaded from http://adsn.ddnetbio.com/. No additional filtering of the data was performed and cell type annotations by Grubman et al. were used. SAVERCAT was run on the dataset using default settings adjusting for three scenarios: (i) age, sex, and library size, (ii) age, sex, PMI, and library size, and (iii) batch and library size. Clustering and UMAP visualization were performed on the original dataset and the SAVERCAT latent representation 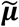 using the standard Seurat workflow. Cells labeled as doublets or unidentified were removed.

Differential expression analysis was performed on the SAVERCAT adjusted counts using the Wilcoxon rank-sum test. Benjamini-Hochberg FDR correction was applied to the raw *p*-values to get the adjusted *p*-values.

## Notes

### Competing Interest Statement

The authors have declared no competing interest.

