## Supplementary figures and images for "Dimension reduction and denoising of single-cell RNA sequencing data in the presence of observed confounding variables"

### Supplementary Fig. 1

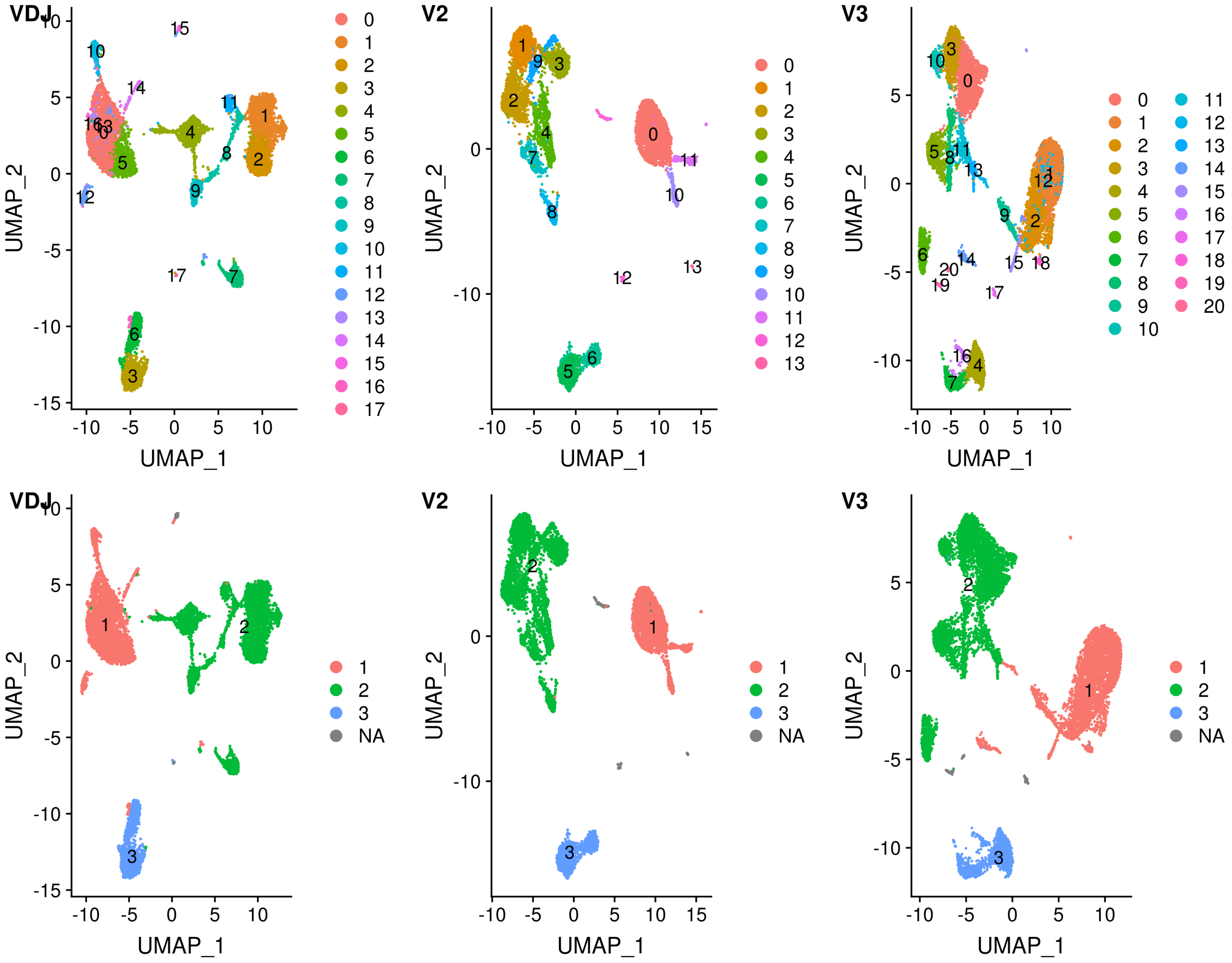

### Supplementary Fig. 2

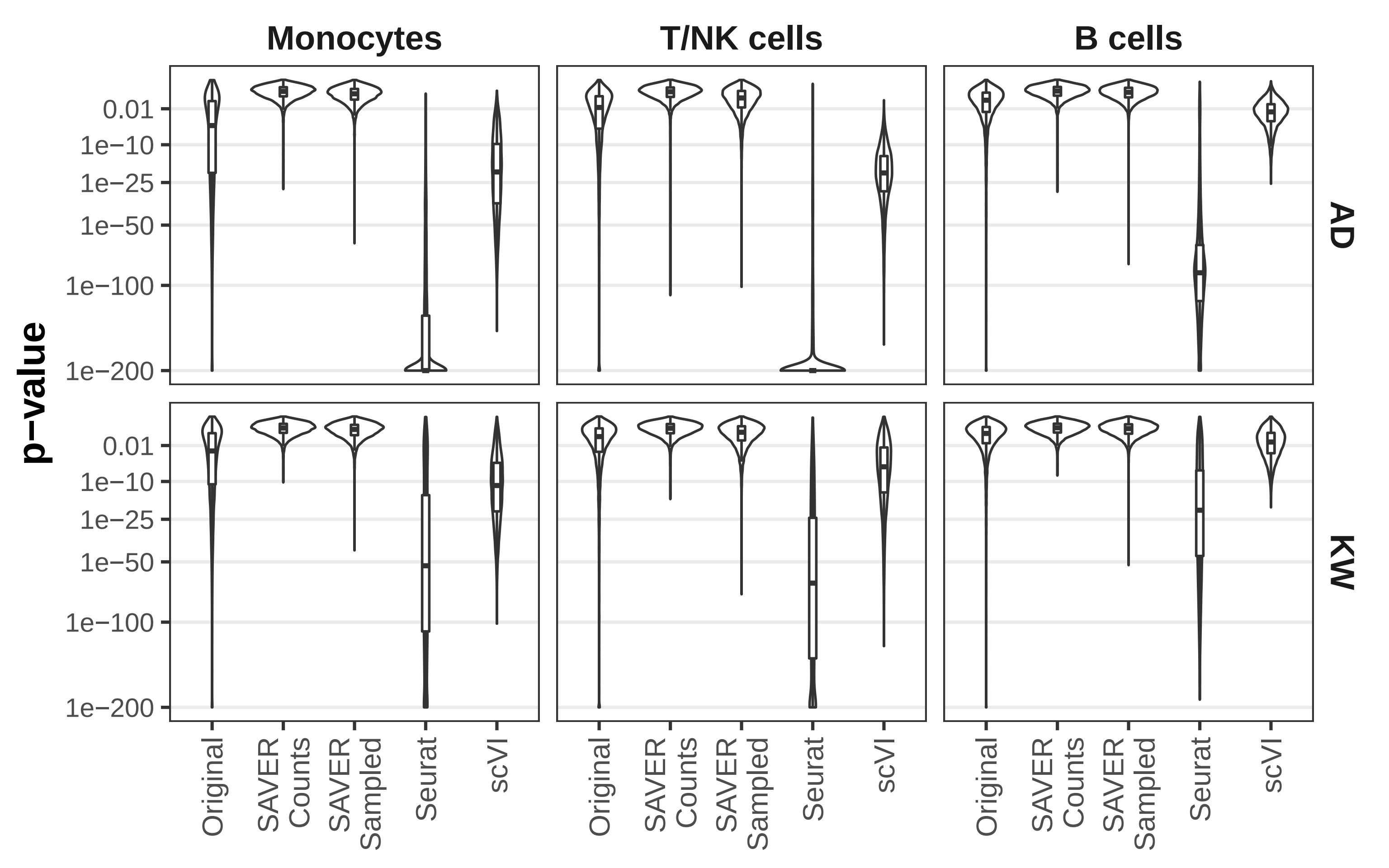

### Supplementary Fig. 3

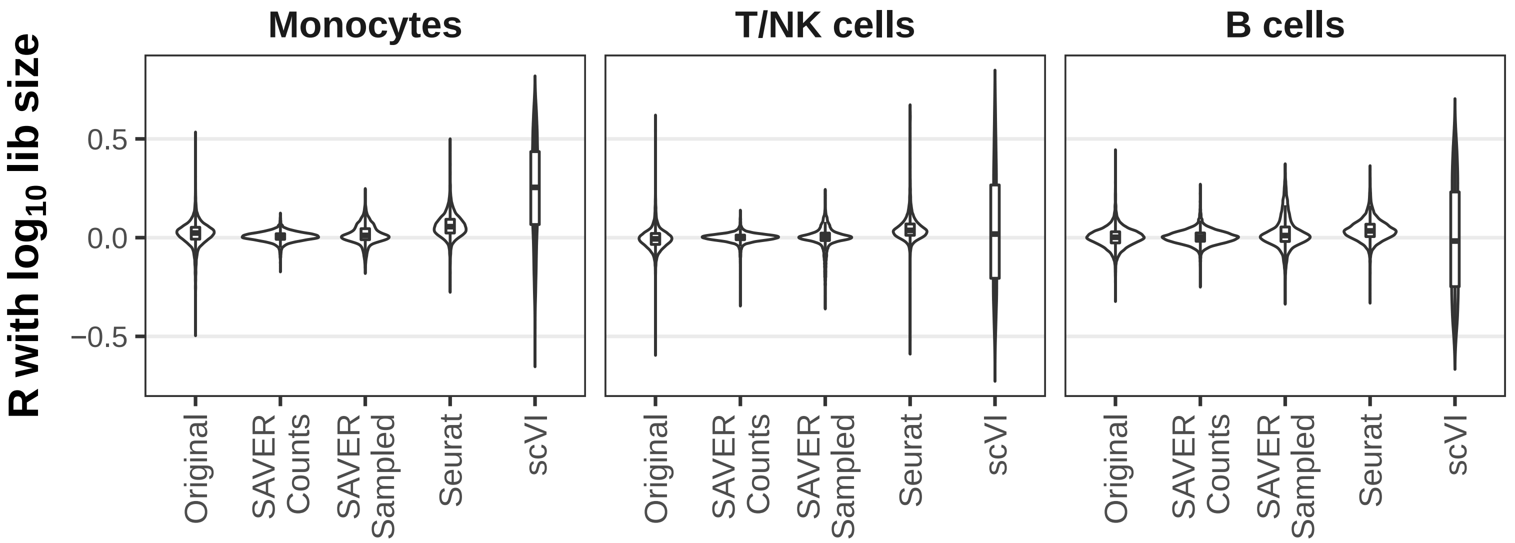

### Supplementary Fig. 4

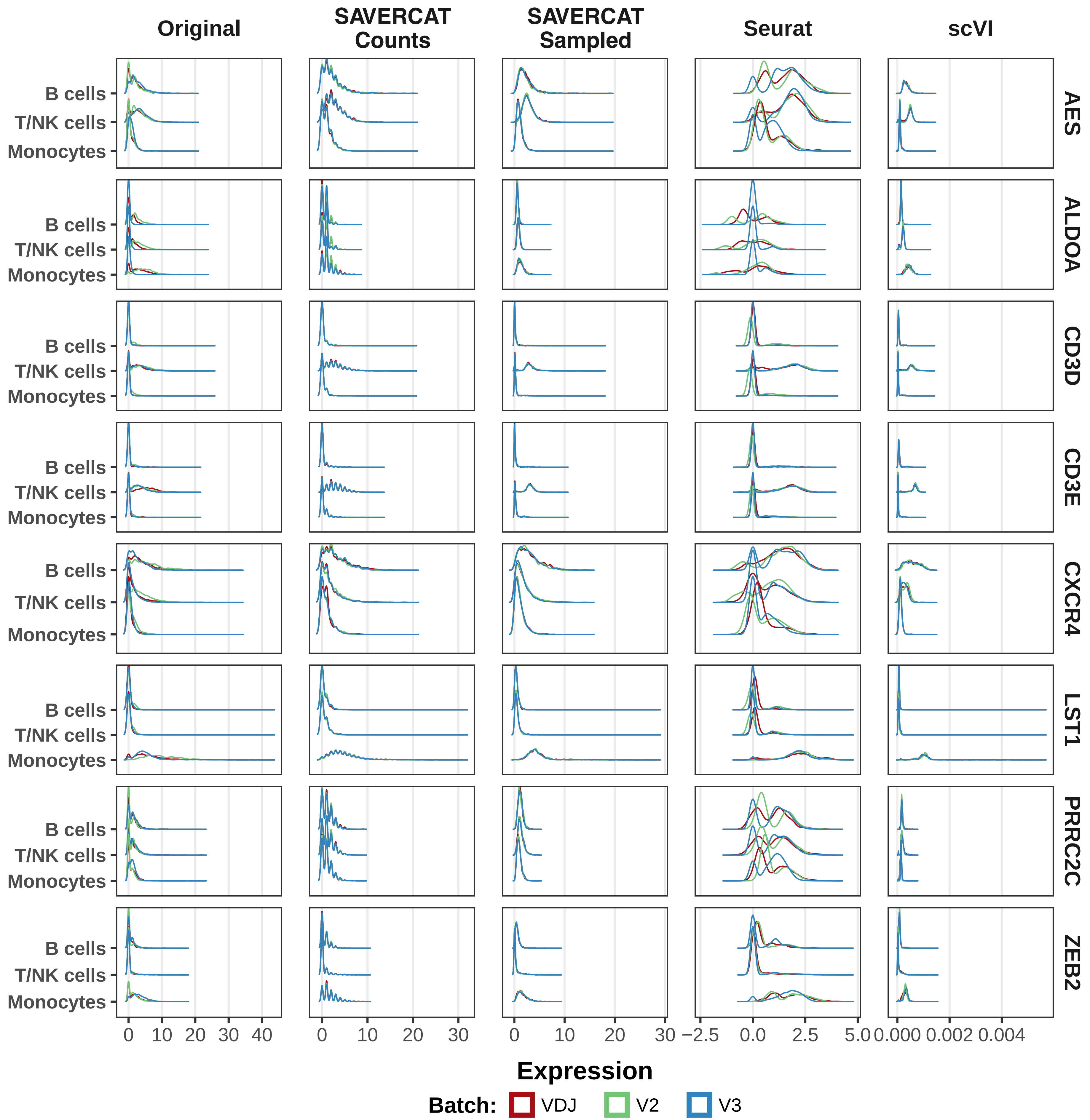

### Supplementary Fig. 5

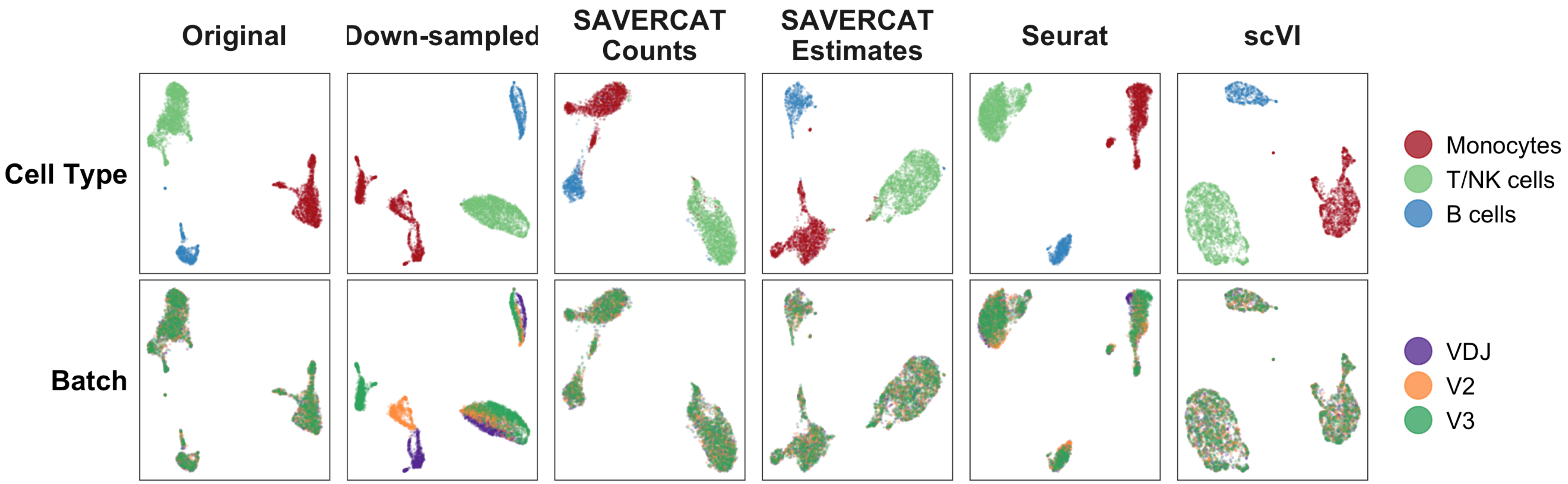

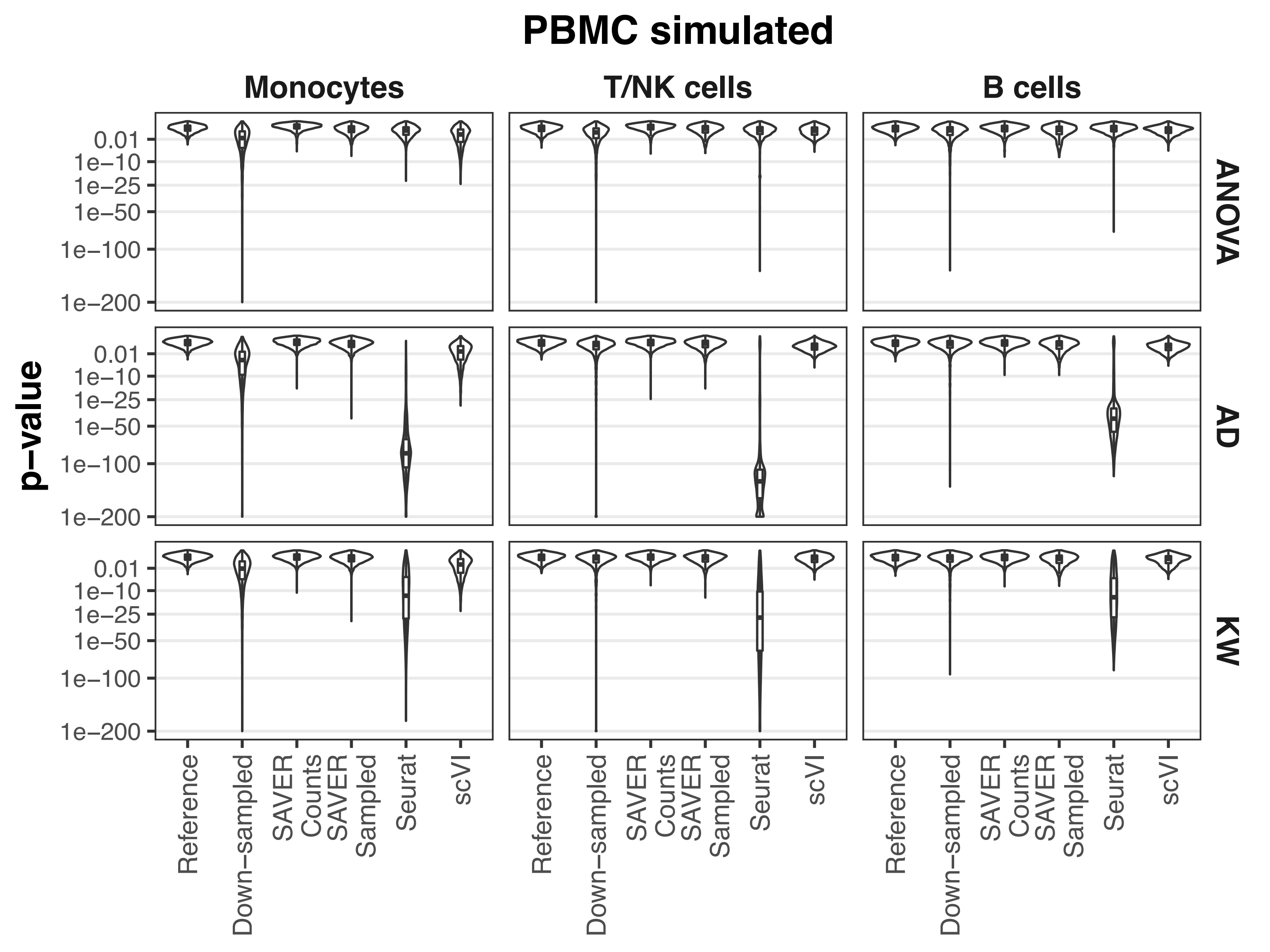

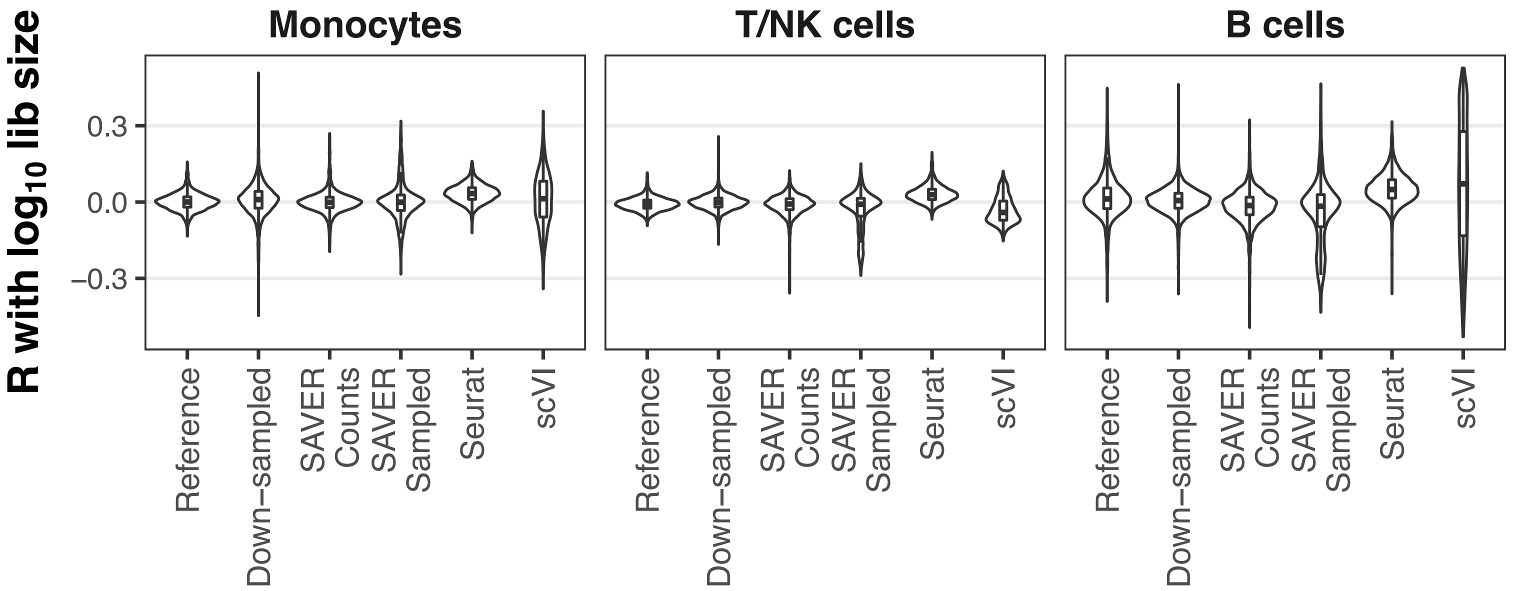

### Supplementary Fig. 6

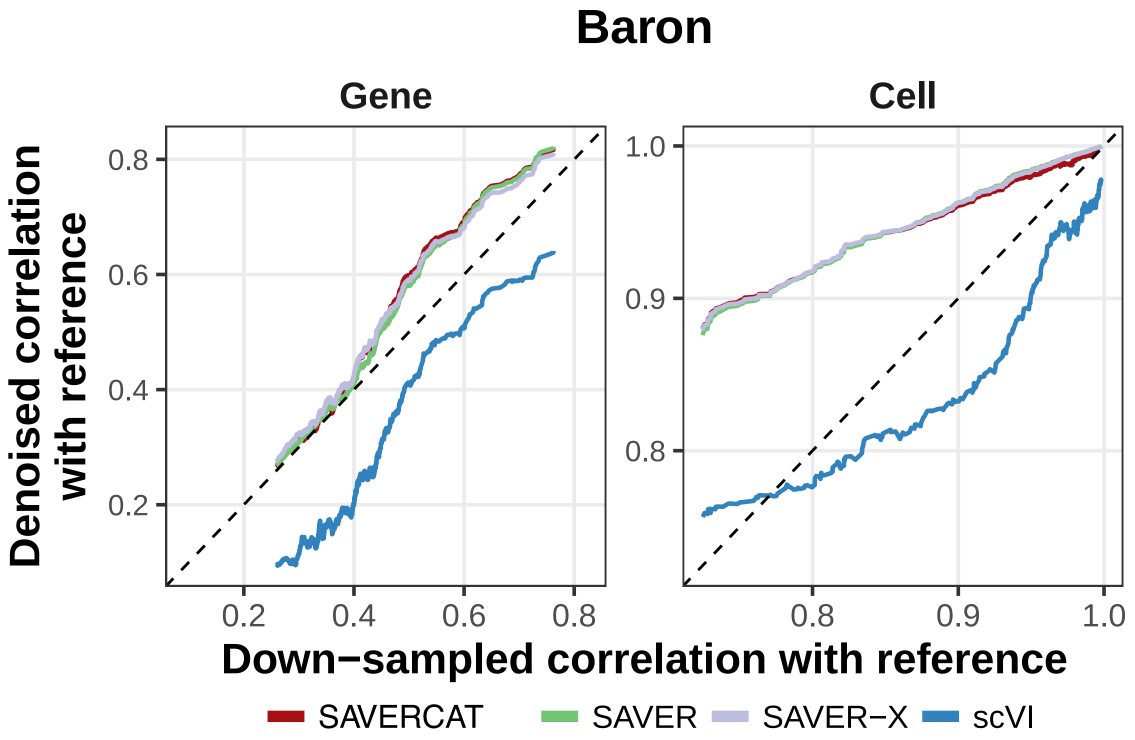

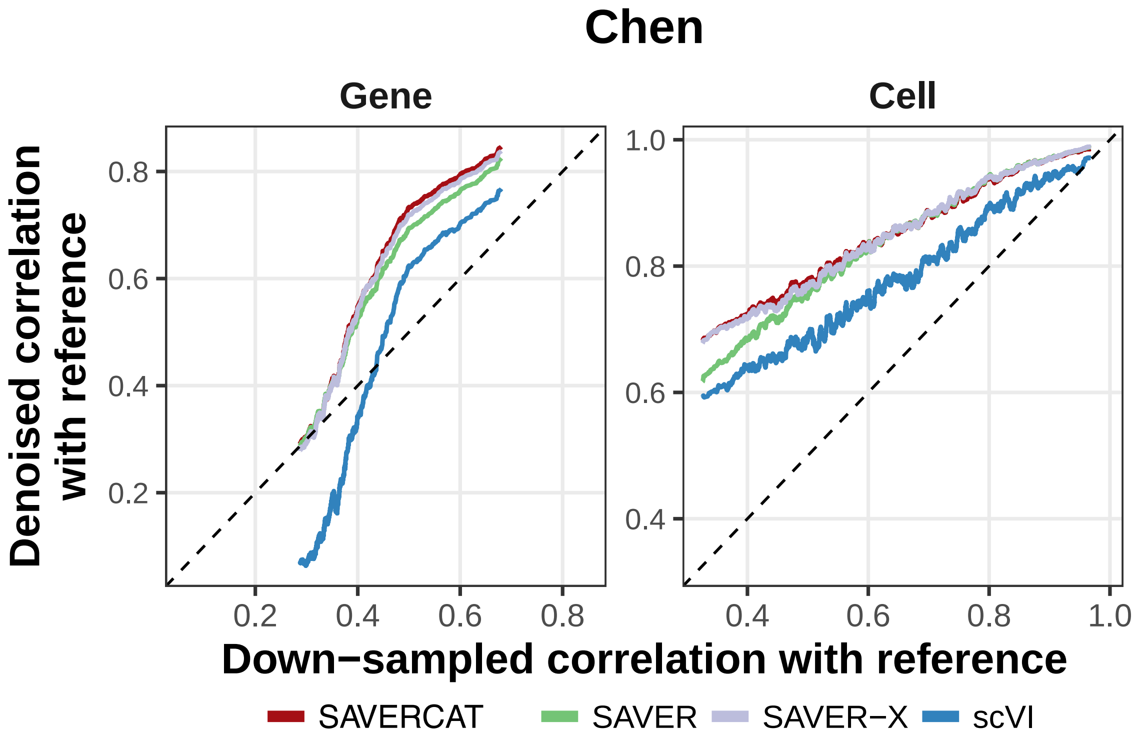

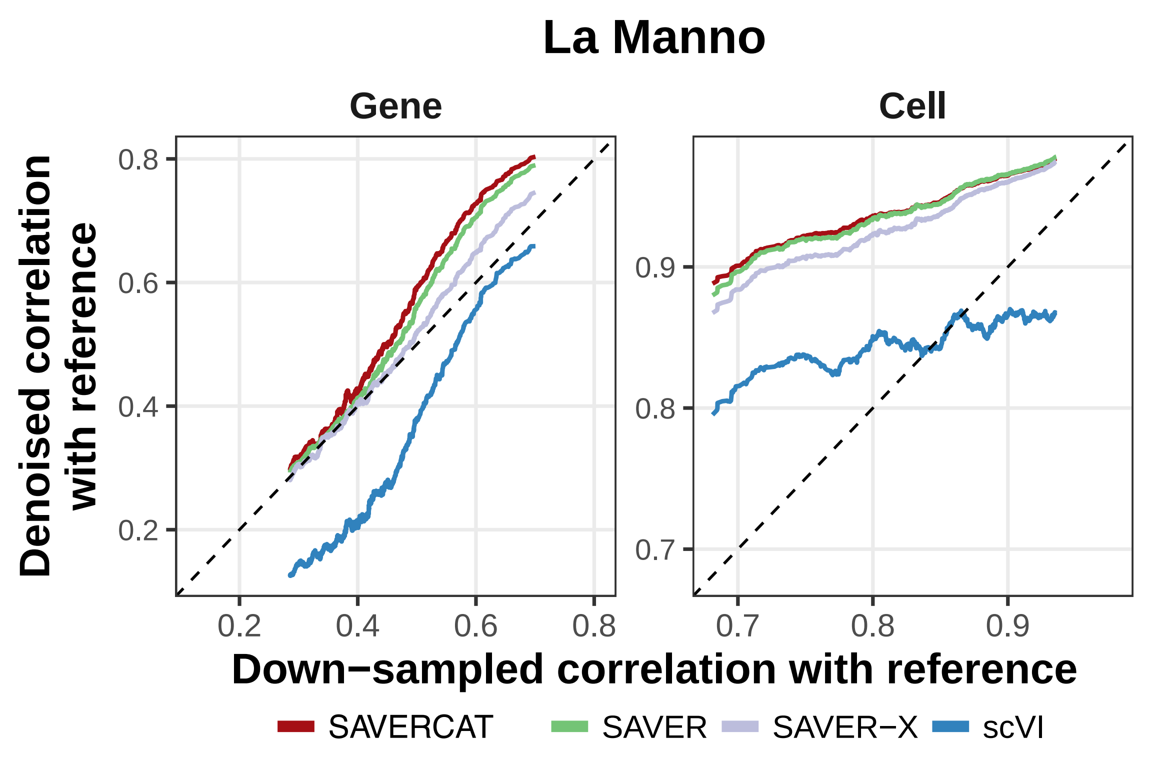

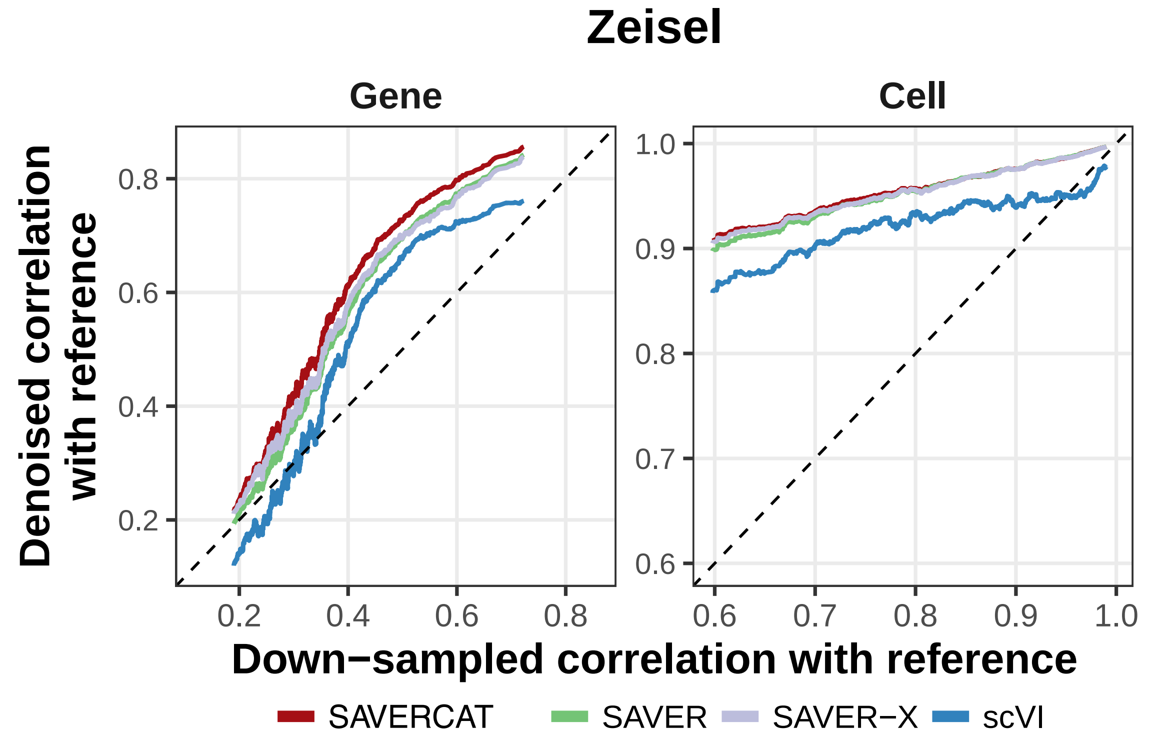

### Supplementary Fig. 7

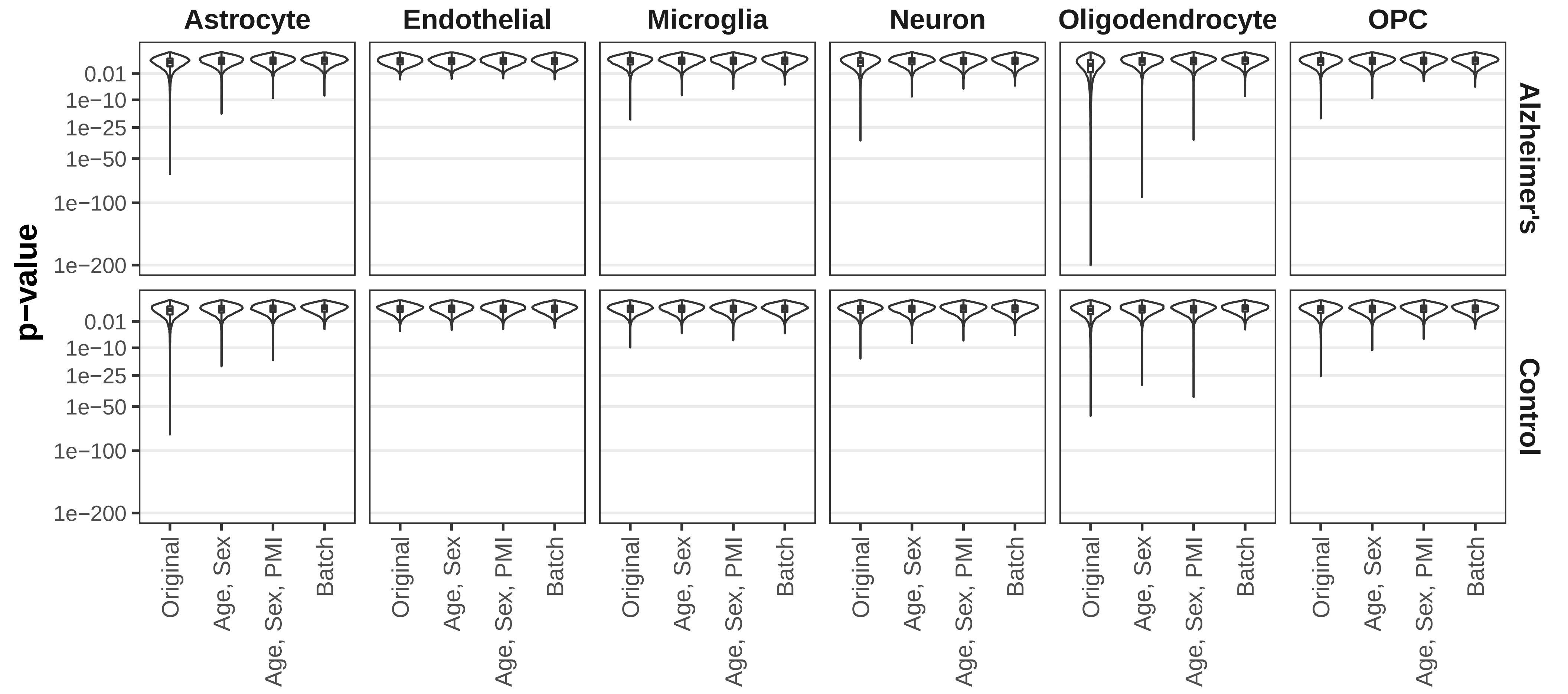
