## Supplementary Table 1 for "Dimension reduction and denoising of single-cell RNA sequencing data in the presence of observed confounding variables"

### PBMC original

### Percentage of genes with p−value < 1e−10

| Celltype | Method | Original | SAVER Counts | SAVER Sampled | Seurat | scVI |
|---|---|---|---|---|---|---|
| Monocytes | ANOVA | 16.7 | 0.0 | 0.0 | 0.4 | 36.2 |
| T/NK cells | ANOVA | 16.0 | 0.0 | 0.5 | 1.5 | 49.9 |
| B cells | ANOVA | 5.5 | 0.0 | 0.0 | 0.5 | 0.8 |
| Monocytes | AD | 38.8 | 0.0 | 0.0 | 98.6 | 74.4 |
| T/NK cells | AD | 16.2 | 0.2 | 1.1 | 99.6 | 88.4 |
| B cells | AD | 6.0 | 0.0 | 0.1 | 96.0 | 3.3 |
| Monocytes | KW | 26.6 | 0.0 | 0.0 | 79.2 | 54.7 |
| T/NK cells | KW | 9.3 | 0.0 | 0.5 | 87.6 | 33.8 |
| B cells | KW | 4.2 | 0.0 | 0.1 | 68.1 | 1.3 |
