## Supplementary Table 2 for "Dimension reduction and denoising of single-cell RNA sequencing data in the presence of observed confounding variables"

PBMC simulated

Percentage of genes with p−value < 1e−10

| Celltype | Method | Reference | Down−sampled | SAVER Counts | SAVER Sampled | Seurat | scVI |
|---|---|---|---|---|---|---|---|
| Monocytes | ANOVA | 0 | 11.4 | 0.0 | 0.0 | 0.1 | 2.3 |
| T/NK cells | ANOVA | 0 | 5.4 | 0.0 | 0.0 | 1.0 | 0.0 |
| B cells | ANOVA | 0 | 2.1 | 0.0 | 0.0 | 0.1 | 0.0 |
| Monocytes | AD | 0 | 23.6 | 0.0 | 0.0 | 98.2 | 4.6 |
| T/NK cells | AD | 0 | 4.6 | 0.1 | 0.1 | 96.9 | 0.0 |
| B cells | AD | 0 | 2.3 | 0.0 | 0.0 | 94.0 | 0.0 |
| Monocytes | KW | 0 | 12.7 | 0.0 | 0.0 | 56.4 | 2.5 |
| T/NK cells | KW | 0 | 3.4 | 0.0 | 0.0 | 75.7 | 0.0 |
| B cells | KW | 0 | 1.8 | 0.0 | 0.0 | 58.6 | 0.0 |
