## Supplementary Table 3 for "Dimension reduction and denoising of single-cell RNA sequencing data in the presence of observed confounding variables"

### Alzheimer's dataset patient information

| Patient | Diagnosis | Batch | Age | Sex | PMI |
|---|---|---|---|---|---|
| AD1 | Alzheimer's | AD1/AD2 | 91.0 | M | 8.0 |
| AD2 | Alzheimer's | AD1/AD2 | 83.8 | M | 10.0 |
| AD3 | Alzheimer's | AD3/AD4 | 67.8 | F | 21.0 |
| AD4 | Alzheimer's | AD3/AD4 | 83.0 | F | 34.0 |
| AD5 | Alzheimer's | AD5/AD6 | 73.0 | M | 9.5 |
| AD6 | Alzheimer's | AD5/AD6 | 74.6 | M | 30.0 |
| Ct1 | Control | Ct1/Ct2 | 67.3 | F | 24.0 |
| Ct2 | Control | Ct1/Ct2 | 82.7 | F | 28.5 |
| Ct3 | Control | Ct3/Ct4 | 72.6 | M | 42.5 |
| Ct4 | Control | Ct3/Ct4 | 75.6 | M | 46.0 |
| Ct5 | Control | Ct5/Ct6 | 77.5 | M | 53.5 |
| Ct6 | Control | Ct5/Ct6 | 82.7 | M | 27.0 |
