## Supplementary Table 4 for "Dimension reduction and denoising of single-cell RNA sequencing data in the presence of observed confounding variables"

Alzheimer's

Percentage of genes with p−value < 1e−10

| Celltype | Diagnosis | Original | Age, Sex | Age, Sex, PMI | Batch |
|---|---|---|---|---|---|
| Astrocyte | Alzheimer's | 0.6 | 0.1 | 0.0 | 0 |
| Endothelial | Alzheimer's | 0.0 | 0.0 | 0.0 | 0 |
| Microglia | Alzheimer's | 0.1 | 0.0 | 0.0 | 0 |
| Neuron | Alzheimer's | 0.2 | 0.0 | 0.0 | 0 |
| Oligodendrocyte | Alzheimer's | 5.7 | 0.2 | 0.0 | 0 |
| OPC | Alzheimer's | 0.1 | 0.0 | 0.0 | 0 |
| Astrocyte | Control | 0.4 | 0.1 | 0.0 | 0 |
| Endothelial | Control | 0.0 | 0.0 | 0.0 | 0 |
| Microglia | Control | 0.0 | 0.0 | 0.0 | 0 |
| Neuron | Control | 0.0 | 0.0 | 0.0 | 0 |
| Oligodendrocyte | Control | 0.6 | 0.1 | 0.1 | 0 |
| OPC | Control | 0.1 | 0.0 | 0.0 | 0 |
