## Supplementary Note 1 for "Dimension reduction and denoising of single-cell RNA sequencing data in the presence of observed confounding variables"

#### Supplementary Note 1: Comparison of 4 loss functions and evaluation of the effect of dropout.

In this note, we consider two factors that may influence the performance of SAVERCAT: The choice of loss function and the utilization of dropout layers.

To define the loss functions, we follow the notation in the main text. Let $s_c$, $\boldsymbol{b_c}$ and $\boldsymbol{y_c}$ denote scaled library size, observed covariate vector and expression level, respectively, of highly variable genes in cell c. Let $\boldsymbol{\widetilde{\mu}_c}, \boldsymbol{\widetilde{\sigma}_c}^{\mathbf{2}}$ be the posterior mean and variance of the normal distribution for the latent variable $\boldsymbol{z}$ estimated by the encoder. Let $\mu'_{cg}$ be the mean of gene g in cell c estimated using the decoder with the input being a sample of $\boldsymbol{z_c}$ concatenated with $\boldsymbol{b_c}$. Let $\mu'_{0_{cg}}$ be the mean from the decoder with the input being the zero vector $\boldsymbol{z}_0 = \mathbf{0}$ concatenated with $\boldsymbol{b_c}$. Let $\theta'_{cg}$ be the dispersion parameter of the negative binomial likelihood, also estimated by the decoder. We considered 4 different loss functions in the first step of training:

(1) NB:

$$\mathcal{L}_\alpha(\boldsymbol{y_c}, \boldsymbol{b_c}; \theta, \phi) = -KL(\boldsymbol{\mathcal{N}}(\boldsymbol{\widetilde{\mu}_c}, \boldsymbol{\widetilde{\sigma}_c}^2\boldsymbol{I})||\boldsymbol{\mathcal{N}}(\mathbf{0}, \boldsymbol{I})) + (1-\alpha)\log\sum_g \mathbf{NegBinom}(Y_{cg}; \mathrm{s_c}\mu'_{cg}, \theta'_{cg}) + \alpha\log\sum_{\mathrm{g}} \mathbf{NegBinom}(Y_{cg}; \mathrm{s_c}\mu'_{0_{cg}}, \theta'_{cg})$$

(2) normNB:

$$\mathcal{L}_\alpha(\boldsymbol{y_c}, \boldsymbol{b_c}; \theta, \phi) = -KL(\boldsymbol{\mathcal{N}}(\boldsymbol{\widetilde{\mu}_c}, \boldsymbol{\widetilde{\sigma}_c}^2\boldsymbol{I})||\boldsymbol{\mathcal{N}}(\mathbf{0}, \boldsymbol{I})) + (1-\alpha)\log\sum_g \mathbf{NegBinom}(Y_{cg}/s_c; \mu'_{cg}, \theta'_{cg}) + \alpha\log\sum_{\mathrm{g}} \mathbf{NegBinom}(Y_{cg}/s_c; \mu'_{0_{cg}}, \theta'_{cg})$$

(3) Poi:

$$\mathcal{L}_\alpha(\boldsymbol{y_c}, \boldsymbol{b_c}; \theta, \phi) = -KL(\boldsymbol{\mathcal{N}}(\boldsymbol{\widetilde{\mu}_c}, \boldsymbol{\widetilde{\sigma}_c}^2\boldsymbol{I})||\boldsymbol{\mathcal{N}}(\mathbf{0}, \boldsymbol{I})) + (1-\alpha)\log\sum_g \mathbf{Poi}(Y_{cg}; \mathrm{s_c}\mu'_{cg}) + \alpha\log\sum_{\mathrm{g}} \mathbf{Poi}(Y_{cg}; \mathrm{s_c}\mu'_{0_{cg}})$$

(4) normPoi:

$$\mathcal{L}_\alpha(\boldsymbol{y_c}, \boldsymbol{b_c}; \theta, \phi) = -KL(\boldsymbol{\mathcal{N}}(\boldsymbol{\widetilde{\mu}_c}, \boldsymbol{\widetilde{\sigma}_c}^2\boldsymbol{I})||\boldsymbol{\mathcal{N}}(\mathbf{0}, \boldsymbol{I})) + (1-\alpha)\log\sum_g \mathbf{Poi}(Y_{cg}/s_c; \mu'_{cg}) + \alpha\log\sum_{\mathrm{g}} \mathbf{Poi}(Y_{cg}/s_c; \mu'_{0_{cg}})$$

Dropout (43) is a widely used regularization method in neural network training. Since a fraction of nodes are randomly zeroed out, dropout has been widely observed to prevent weights for different nodes from aligning and avoid overfitting to some outlier data points. In this experiment, we consider adding a dropout layer to in the first encoder layer with a dropout fraction of 0.1.

We applied SAVERCAT using each of the four loss functions above with or without dropout to benchmarking dataset 4 (Human Pancreas data), 5 (PBMC data) and 6 (Cell Line data) from Tran et al (37), and evaluated the effects by visualization using t-SNE and UMAP and calculating LISI scores.

### 1. Dataset 5

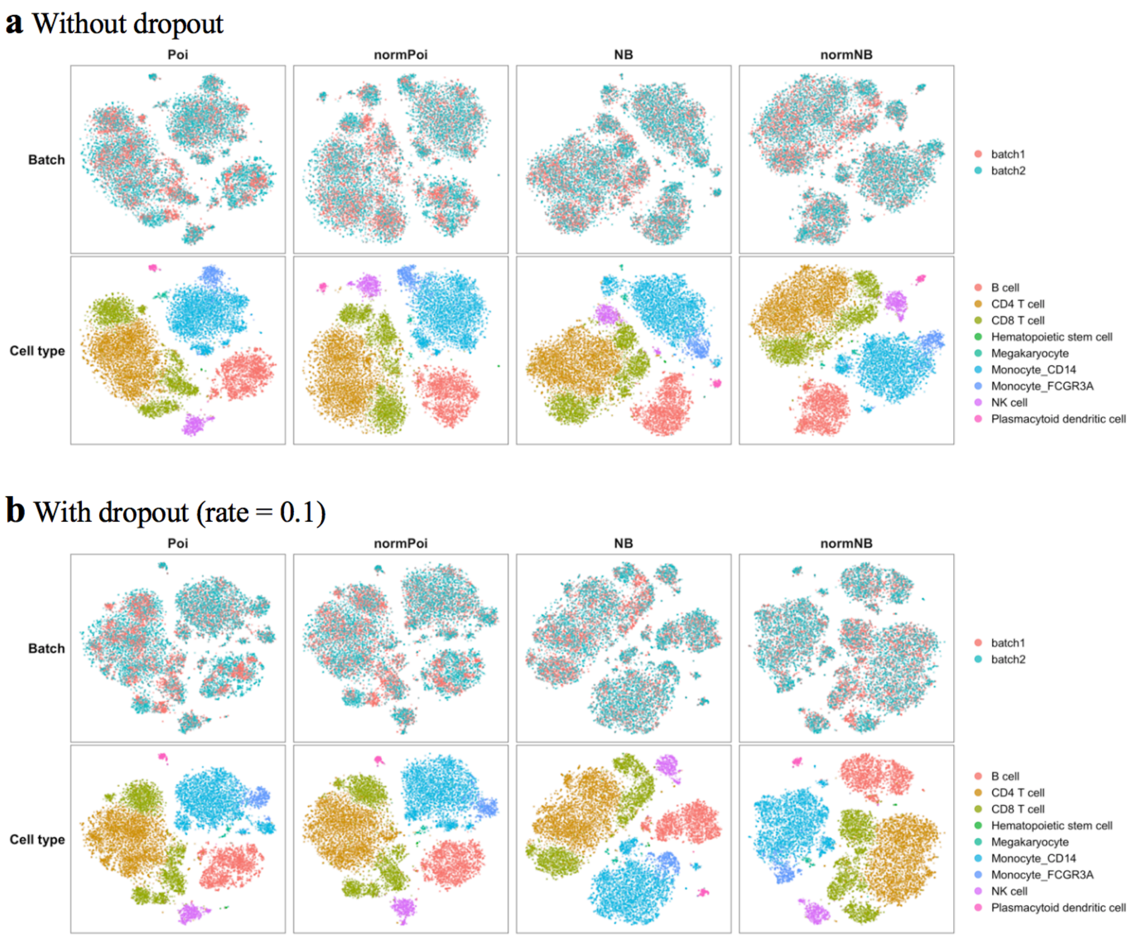


**Figure 1** t-SNE visualization of the performance of SAVERCAT using 4 different loss functions with (**a**) or without (**b**) dropout on benchmarking dataset 5. $\alpha$ in loss function is set to 0.01. Highly variable genes are chosen across the entire dataset and dispersion parameter of negative binomial distribution is defined as cell and gene specific. The other running options are set as default.

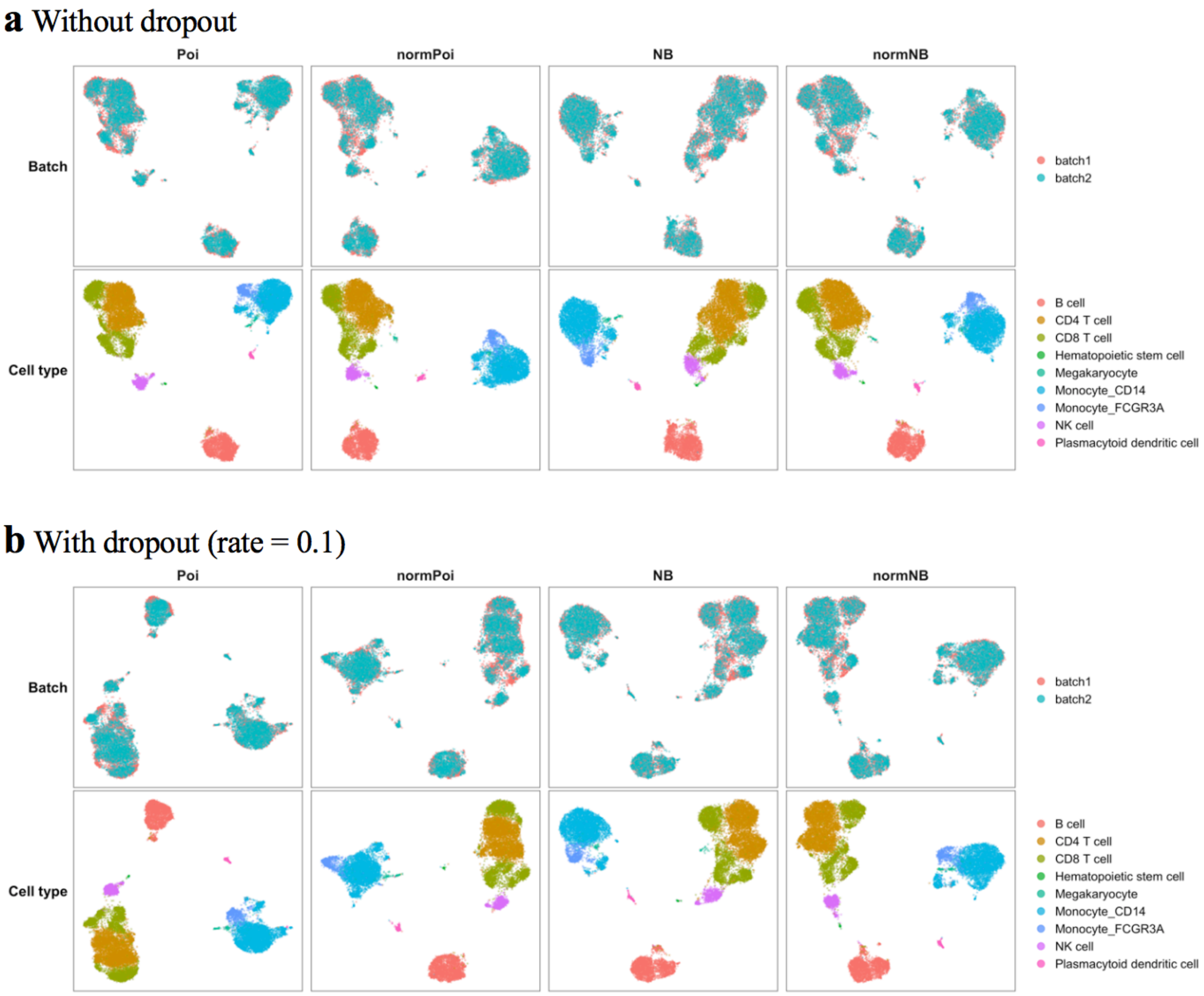


**Figure 2** UMAP visualization of the performance of SAVERCAT using 4 different loss functions with (**a**) or without (**b**) dropout on benchmarking dataset 5. $\alpha$ in loss function is set to 0.01. Highly variable genes are chosen across the entire dataset and dispersion parameter of negative binomial distribution is defined as cell and gene specific. The other running options are set as default.

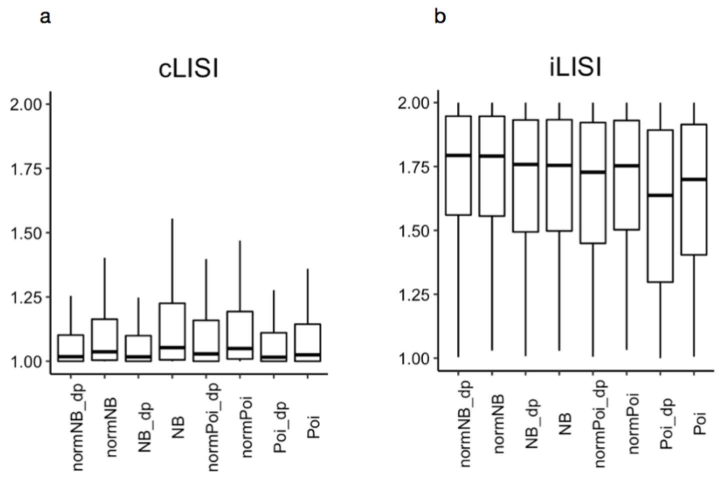


**Figure 3** LISI scores for evaluating the performance of SAVERCAT using 4 different loss functions with or without dropout on benchmarking dataset 5. "dp" means using dropout. Both cLISI (a) and iLISI (b) are unscaled results calculated using the *lisi* R package with default perplexity=30.

For loss function using Poisson likelihood, there are tiny clusters within batch 1 in t-SNE plot (Fig. 1 **a**-Poi, **b**-Poi). When we use negative binomial loss, tiny clusters are less obvious and batches are better mixed (Fig. 1 **a**-NB, **b**-NB). This phenomenon is also observed when dropout is used (Fig. 1 **b**). A possible interpretation is that the outlying highly dispersed genes are weighted more in Poisson loss than in negative binomial loss. Normalization before calculating likelihood (Fig. 1 **a**-normPoi, **a**-normNB, **b**-normPoi, **b**-normNB) will make points more scattered and lead to better batch mixing. In this dataset, CD4 T cell and CD8 T cell as well as CD14 Monocyte cell and FCGR3A Monocyte cell are 2 pairs of similar cells. In both t-SNE and UMAP visualization, SAVERCAT with dropout (Fig. 1 **b**, Fig. 2 **b**) does better across 4 loss functions in that it separates CD4 T cells from CD8 T cells better and cleans up isolated points on the edges of clusters to make the cluster cleaner and tighter.

#### 2. Dataset 6

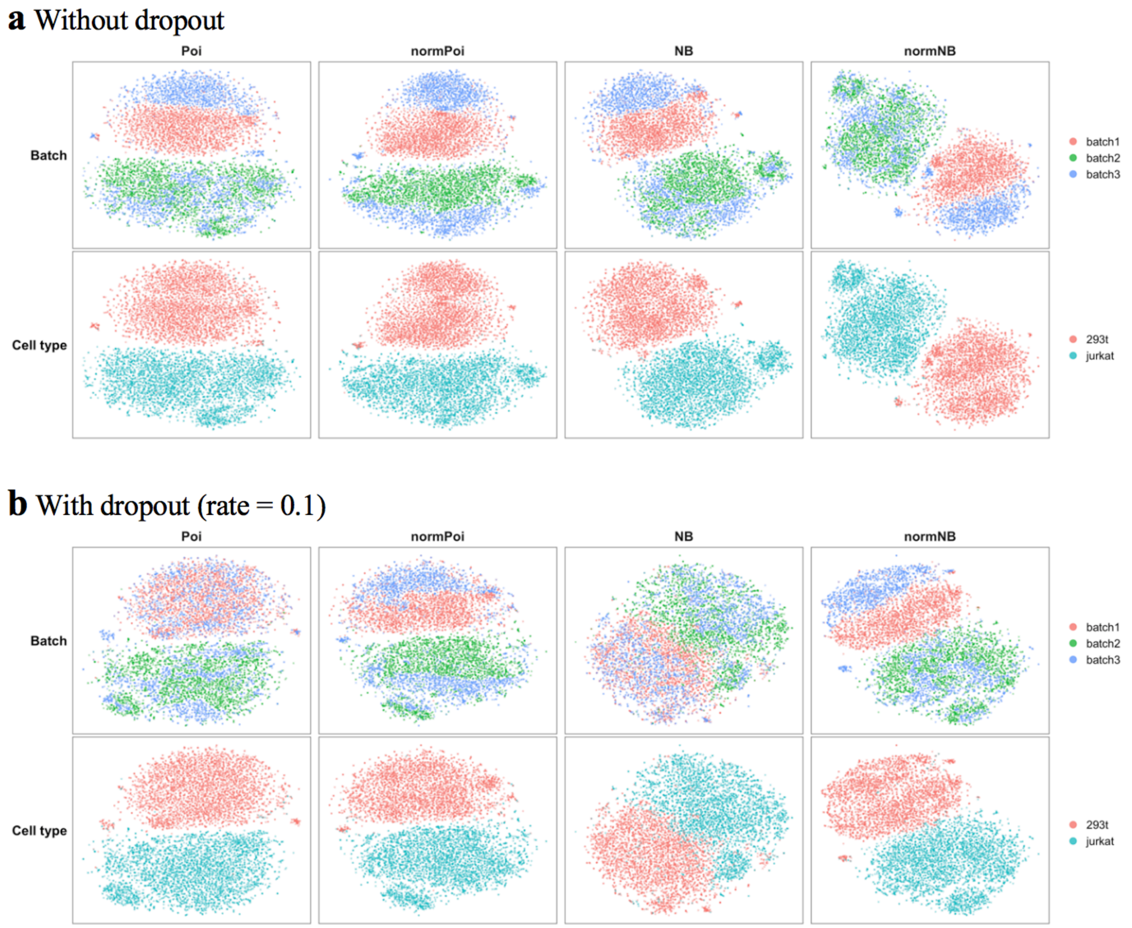


**Figure 4** t-SNE visualization of the performance of SAVERCAT using 4 different loss functions with (**a**) or without (**b**) dropout on benchmarking dataset 6. $\alpha$ in loss function is set to 0.01. Highly variable genes are chosen across the entire dataset and dispersion parameter of negative binomial distribution is defined as cell and gene specific. The other running options are set as default.

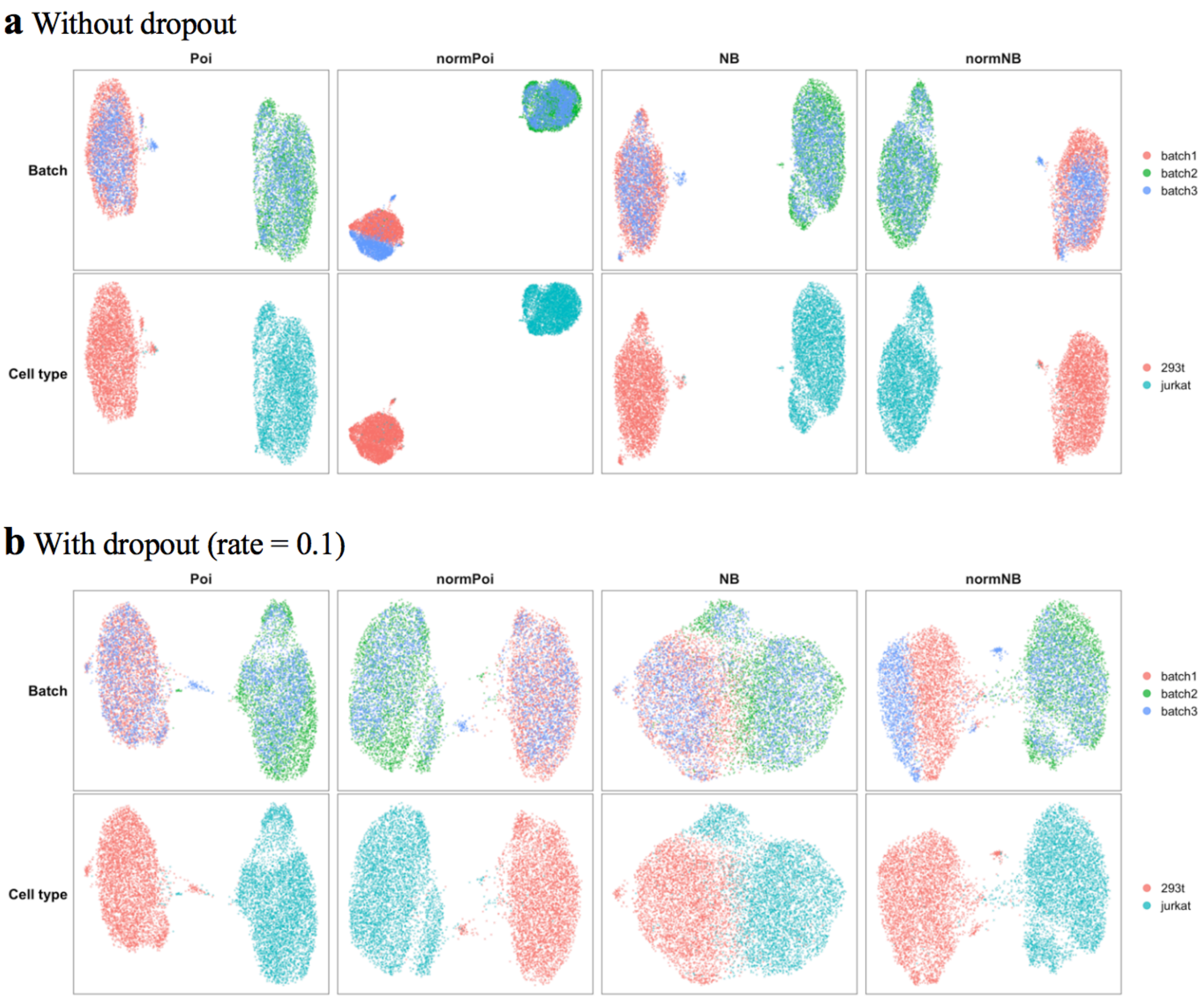


**Figure 5** UMAP visualization of the performance of SAVERCAT using 4 different loss functions with (**a**) or without (**b**) dropout on benchmarking dataset 6. $\alpha$ in loss function is set to 0.01. Highly variable genes are chosen across the entire dataset and dispersion parameter of negative binomial distribution is defined as cell and gene specific. The other running options are set as default.

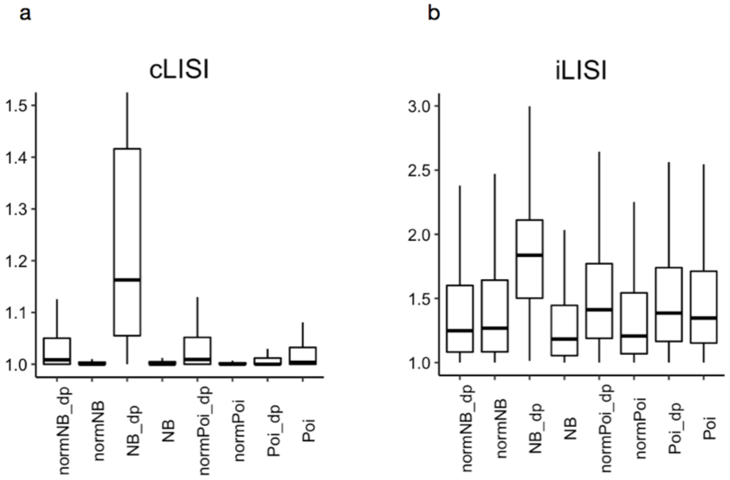


**Figure 6** LISI scores for evaluating the performance of SAVERCAT using 4 different loss functions with or without dropout on benchmarking dataset 6. "dp" means using dropout. Both cLISI (a) and iLISI (b) are unscaled results calculated using the *lisi* R package with default perplexity=30.

For dataset 6, in the t-SNE plot (Fig. 4), SAVERCAT with dropout works better in terms of mixing 293t cells from batch 1 and batch 3. When using dropout, both SAVERCAT with negative binomial loss and SAVERCAT with Poisson loss mix batches well. However, it is noticed from both t-SNE (Fig. 4) and UMAP plot (Fig. 5) that SAVERCAT with negative binomial loss makes 293T and Jurkat cells, which are significantly different cell types, mix a little. The underlying reason might be SAVERCAT using negative binomial loss estimates cell specific dispersion parameter, so that it absorb part of characterizations of cells into the dispersion parameter instead of the latent means which are used in visualization. This also leads us to think of modeling batch specific dispersion parameters separately from modeling negative binomial mean, which is the default option of our final model described in detail in the main paper. From Fig. 6, there is a positive correlation between iLISI and cLISI, which indicates methods separating cell types better may do worse in batch mixing.

#### 3. Dataset 4

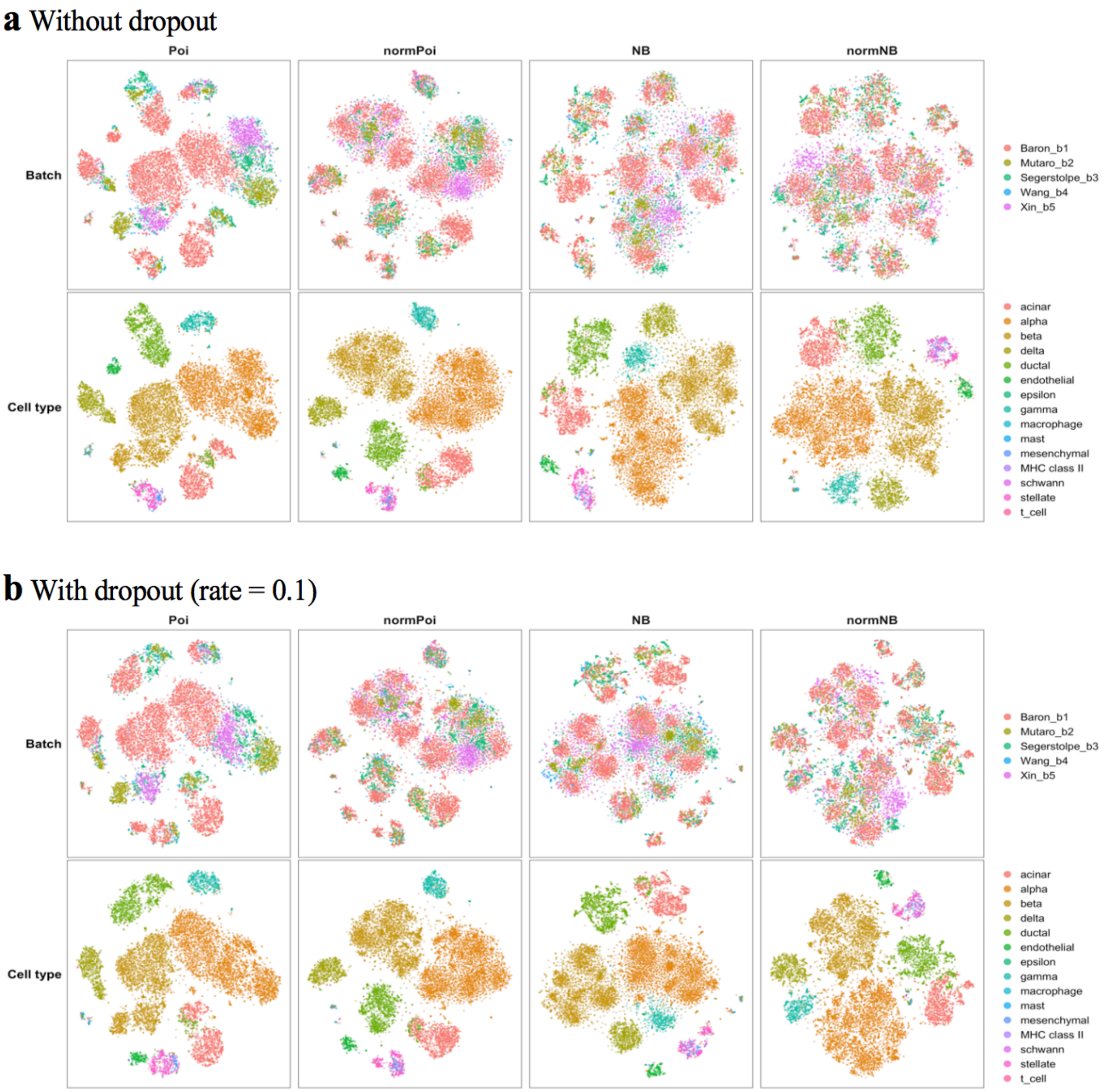


**Figure 7** t-SNE visualization of the performance of SAVERCAT using 4 different loss functions with (**a**) or without (**b**) dropout on benchmarking dataset 4. $\alpha$ in loss function is set to 0.01. Highly variable genes are chosen across the entire dataset and dispersion parameter of negative binomial distribution is defined as cell and gene specific. The other running options are set as default.

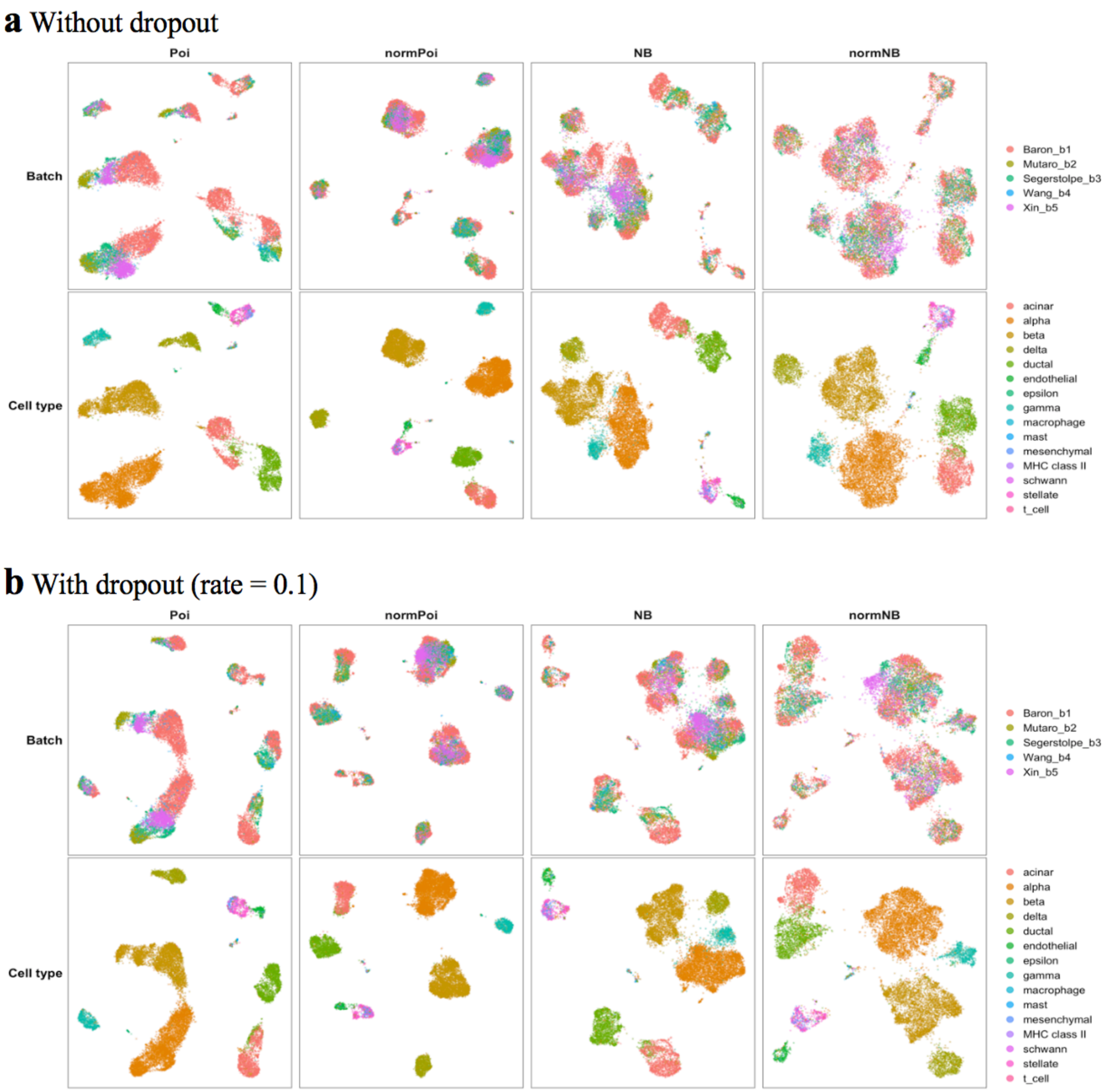


**Figure 8** UMAP visualization of the performance of SAVERCAT using 4 different loss functions with (**a**) or without (**b**) dropout on benchmarking dataset 4. $\alpha$ in loss function is set to 0.01. Highly variable genes are chosen across the entire dataset and dispersion parameter of negative binomial distribution is defined as cell and gene specific. The other running options are set as default.

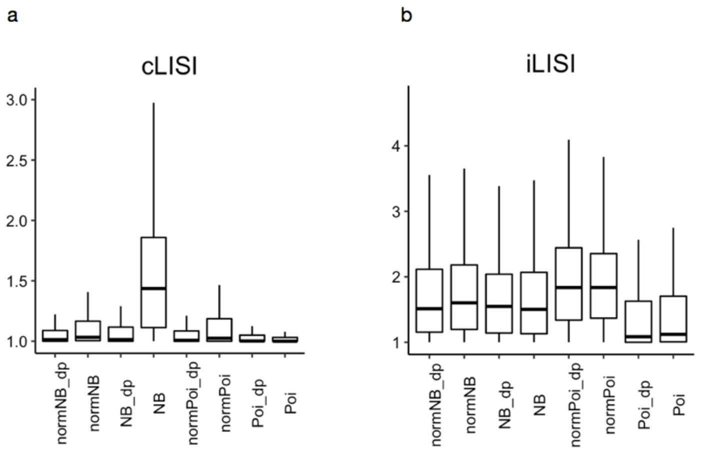


**Figure 9** LISI scores for evaluating the performance of SAVERCAT using 4 different loss functions with or without dropout on benchmarking dataset 4. "dp" means using dropout. Both cLISI (a) and iLISI (b) are unscaled results calculated using the *lisi* R package with default perplexity=30.

From t-SNE plot (Fig. 7), using dropout can make the cluster tighter and clean up isolated points around clusters. Normalization and Poisson performs best in terms of iLISI and cLISI (Fig. 9). We can also see from the t-SNE plot and UMAP plot (Fig. 7, Fig. 8) that compared to Poisson loss, negative binomial loss makes the points more scattered in a uniform way across batches. Normalization also makes points scattered.

#### 4. Conclusions

To summarize, negative binomial loss makes points more scattered which may help in batch mixing. SAVERCAT using Poisson loss is sensitive to outliers and may not be able to distinguish between similar cell types well. The effect of using normalization before calculating likelihood is not obvious. It might have a slight influence on making points more scattered. Using dropout can help in batch mixing and produces cleaner and tighter cell type clusters.
