## Supplementary Note 2 for "Dimension reduction and denoising of single-cell RNA sequencing data in the presence of observed confounding variables"

### Supplementary Note 2: Comparison of different strategies for choosing highly variable genes and estimating dispersion parameters

In this note, we consider two aspects of our method: The method for choosing highly variable genes and the model for the dispersion parameters of the negative binomial likelihood.

In the first step of training, only highly variable genes are used to train the variational autoencoder. How we select highly variable genes may make a big difference on the trained model. We considered 2 strategies for choosing them. The first strategy, denoted by "bhvg", calculates highly variable genes for each batch, takes the union across batches, and selects the top 3000 genes from this union. The second strategy is called "chvg", which calculates 3000 highly variable genes across the complete dataset. We tested these 2 strategies by applying SAVERCAT to benchmarking dataset 4 (Human Pancreas data), 5 (PBMC data) and 6 (Cell Line data) from Tran et al, and evaluated the effects by visualization using t-SNE and UMAP and LISI scores.

In supplementary note 1, SAVERCAT with negative binomial loss mixes distinct cell types (293t and Jurcat) in benchmarking dataset 6. It is possibly because dispersion parameters of negative binomial loss are defined as gene and cell specific which absorb some intrinsic cell type differences. This method, which we denote by "cdisp", estimates dispersion parameters together with the means of the negative binomial distribution using the decoder. To improve cell type identification in the latent space, we introduce another model called "bdisp" for the dispersion parameters of the negative binomial loss. This model assumes that the batch specific dispersion parameters rely only on the covariate vector $\boldsymbol{b_c}$. To be specific, we use a decoder to model the negative binomial means as a function of $\boldsymbol{b_c}$ and latent variable $\boldsymbol{z_c}$, and another decoder taking only $\boldsymbol{b_c}$ as input to model the dispersion parameters. We investigate the effectiveness of these two different dispersion models by applying SAVERCAT to benchmarking dataset 4 (Human Pancreas data), 5 (PBMC data) and 6 (Cell Line data) from Tran et al (37) and training it using batch specific highly variable genes. t-SNE plot, UMAP plot and LISI are used for evaluation.

In the training, NB and normNB are 2 loss functions which calculate likelihood based on negative binomial distribution. The difference is normNB uses the scaled counts and NB uses the observed counts. Dropout is used in the first layer of encoder with a dropout rate of 0.1.

### 1. Dataset 5

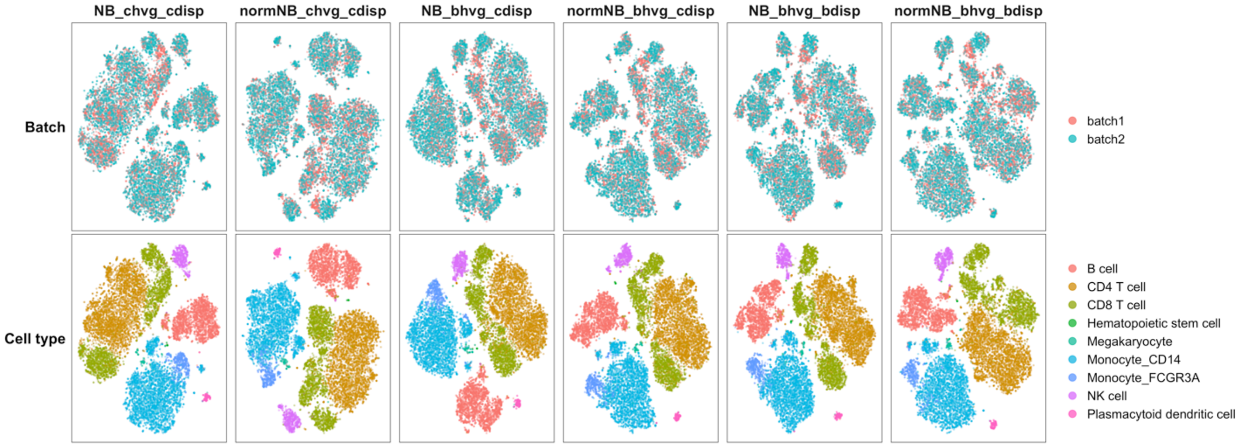


**Figure 1** t-SNE visualization of the performance of SAVERCAT trained with different highly variable genes and different types of dispersion parameters on benchmarking dataset 5.

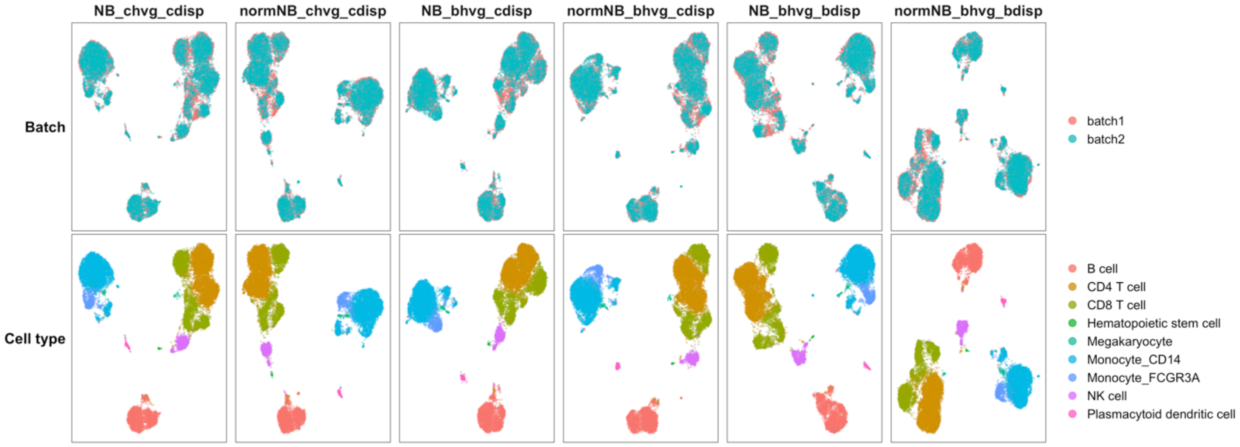


**Figure 2** UMAP visualization of the performance of SAVERCAT trained with different highly variable genes and different types of dispersion parameters on benchmarking dataset 5.

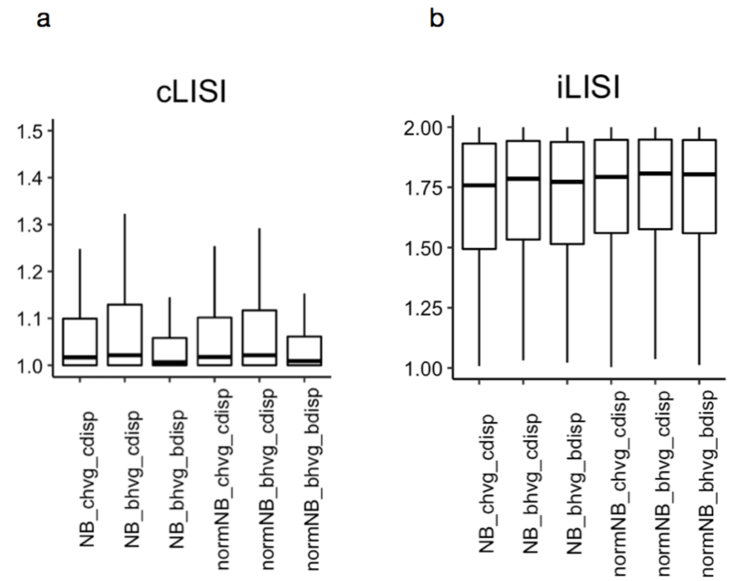


**Figure 3** LISI scores for evaluating the performance of SAVERCAT trained with different highly variable genes and different types of dispersion parameters on benchmarking dataset 5. Both cLISI (a) and iLISI (b) are unscaled results calculated using the *lisi* R package with default perplexity=30.

SAVERCAT with batch specific dispersion parameters does better than with cell specific dispersion parameters in terms of separating different cell types. For SAVERCAT using normalization negative binomial loss with batch specific dispersion parameters which is trained using concatenated batch specific highly variable genes (normNB_bhvg_bdisp in Fig. 1 and Fig. 2), CD8 T cells are well separated from CD4 T cells, and even 2 types of Monocyte cells which are very similar can be separated. cLISI scores (Fig. 3a) also indicate that using batch specific dispersion parameters can improve the performance of SAVERCAT in cell type identification.

### 2. Dataset 6

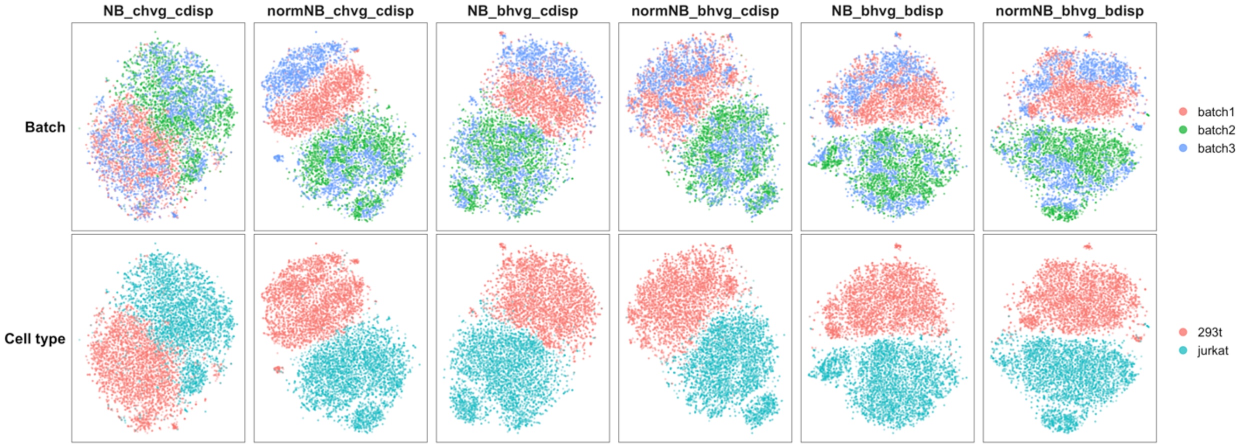


**Figure 4** t-SNE visualization of the performance of SAVERCAT trained with different highly variable genes and different types of dispersion parameters on benchmarking dataset 6.

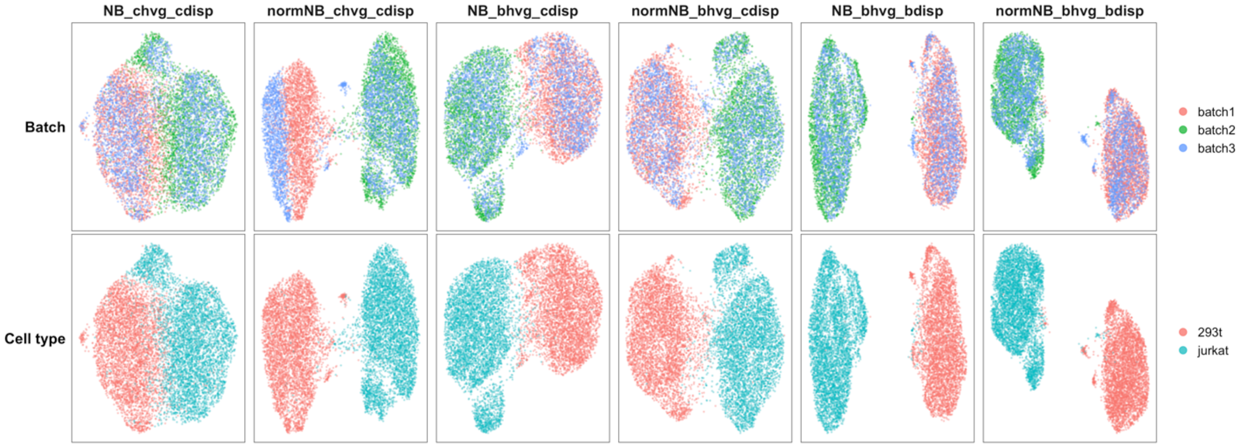


**Figure 5** UMAP visualization of the performance of SAVERCAT trained with different highly variable genes and different types of dispersion parameters on benchmarking dataset 6.

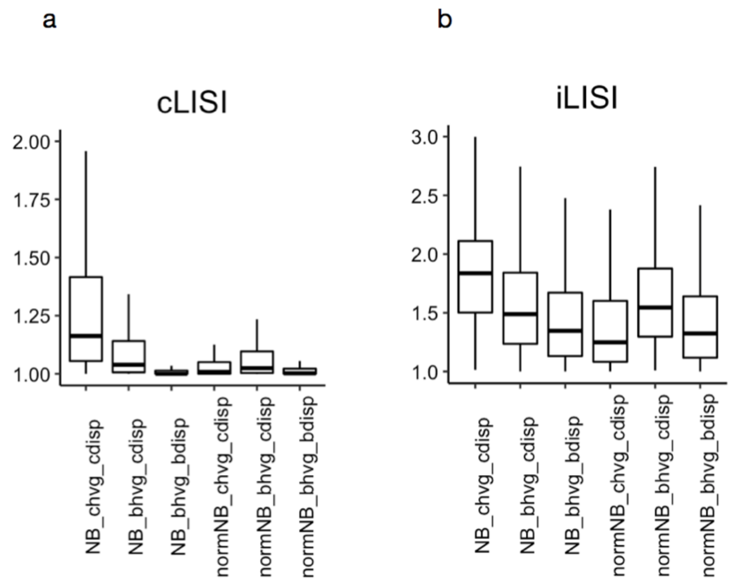


**Figure 6** LISI scores for evaluating the performance of SAVERCAT trained with different highly variable genes and different types of dispersion parameters on benchmarking dataset 6. Both cLISI (a) and iLISI (b) are unscaled results calculated using the *lisi* R package with default perplexity=30.

From Fig. 4, 5, 6, SAVERCAT using negative binomial loss or normalization negative binomial loss with batch specific dispersion parameters and batch specific highly variable genes does the best in terms of separating 2 distinct cell types. Using batch specific highly variable genes can avoid cell type mixing by absorbing cell differences in mean parameter instead of dispersion parameter. It is observed from iLISI and cLISI scores (Fig. 6) that there is a trade off between cell type separation and batch mixing. For example, SAVERCAT using negative binomial loss with cell specific dispersion parameters and complete highly variable genes (NB_chvg_cdisp in Fig. 4, 5, 6) does the best in batch mixing but worst in separating 293t cells from jurcat cells.

### 3. Dataset 4

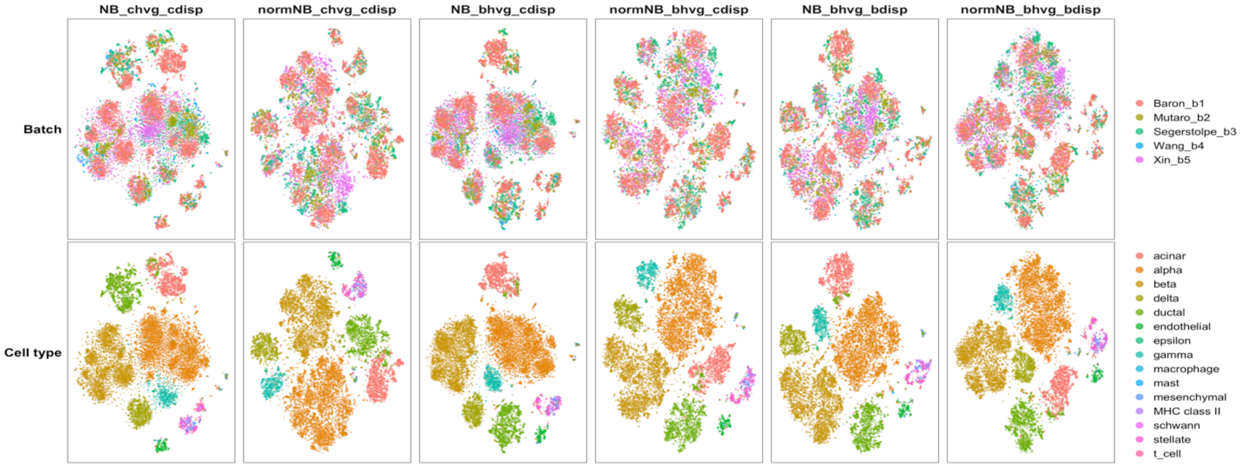


**Figure 7** t-SNE visualization of the performance of SAVERCAT trained with different highly variable genes and different types of dispersion parameters on benchmarking dataset 4.

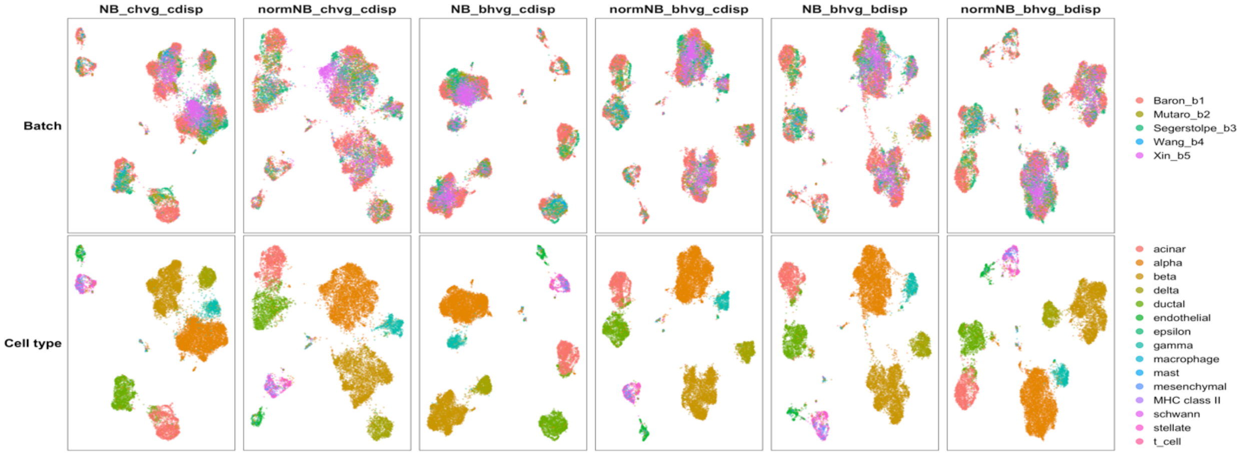


**Figure 8** UMAP visualization of the performance of SAVERCAT trained with different highly variable genes and different types of dispersion parameters on benchmarking dataset 4.

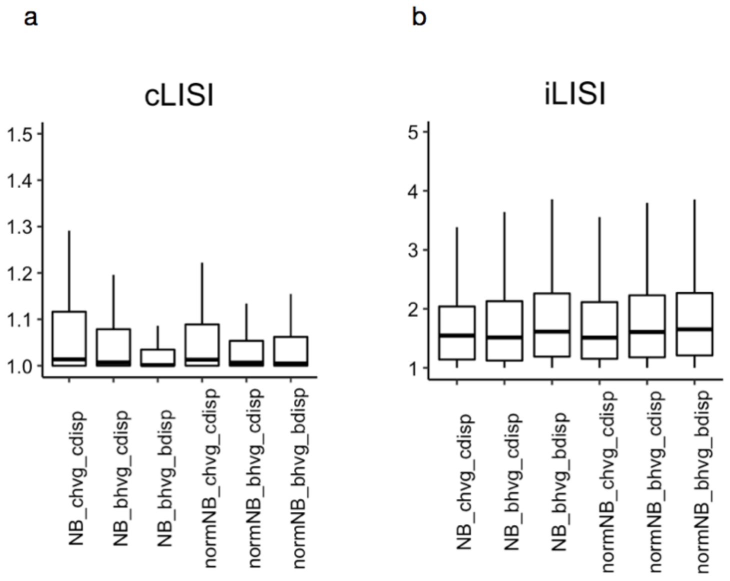


**Figure 9** LISI scores for evaluating the performance of SAVERCAT trained with different highly variable genes and different types of dispersion parameters on benchmarking dataset 4. Both cLISI (a) and iLISI (b) are unscaled results calculated using the *lisi* R package with default perplexity=30.

From Fig. 7, 8, 9, comparing SAVERCAT with complete highly variable genes (NB_chvg_cdisp and normNB_chvg_cdisp) to SAVERCAT with concatenated batch specific highly variable genes (NB_bhvg_cdisp and normNB_bhvg_cdisp), the latter one produces tighter and cleaner clusters and has lower cLISI scores which indicates using batch specific highly variable genes can help in cell type identification. It is shown in Fig. 9 that using batch specific dispersion parameters can improve the performance of SAVERCAT in both distinguishing among cell types and batch mixing.

### 4. Conclusions

To sum up, using batch specific dispersion parameters can avoid cell type mixing by absorbing cell differences into mean parameter instead of dispersion parameter. Using concatenated batch specific highly variable genes can help identifying cell types when differences among cell types are significant like the case of dataset 6 and 4.
