## Supplementary Note 3 for "Dimension reduction and denoising of single-cell RNA sequencing data in the presence of observed confounding variables"

### Supplementary Note 3: UMAP and t-SNE visualization of SAVERCAT, Seurat, scVI and Harmony on benchmarking datasets

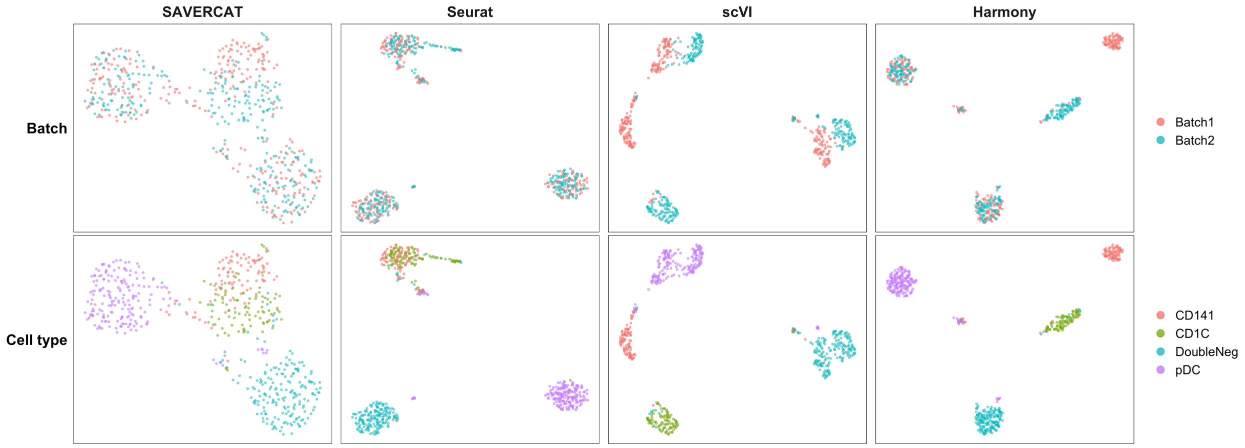


**Figure 1** UMAP visualization of 576 non-UMI human dendritic cells from 2 batches (dataset 1) to evaluate 4 methods: SAVERCAT, Seurat, scVI and Harmony.

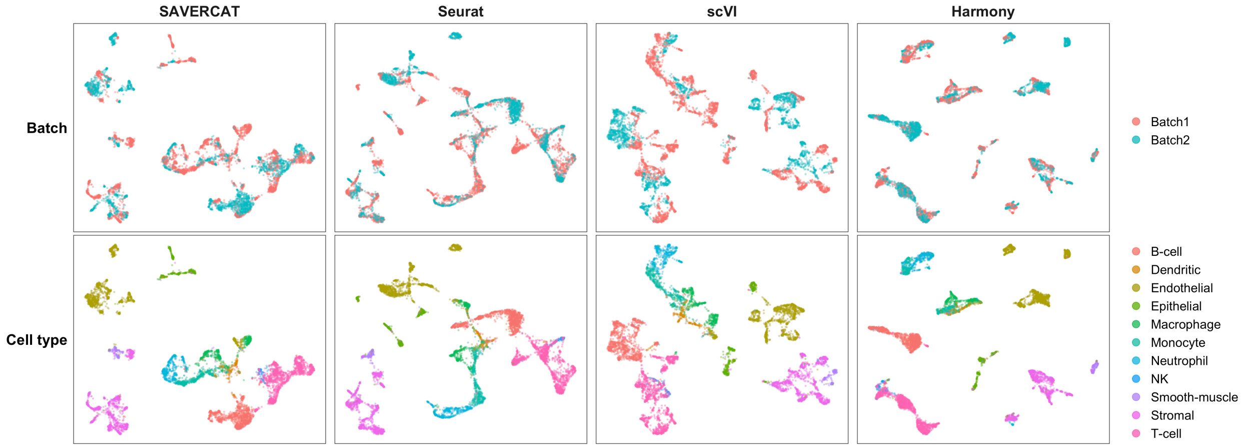


**Figure 2** UMAP visualization of 6,954 non-UMI cells from two Mouse Cell Atlas studies (dataset 2) to evaluate 4 methods: SAVERCAT, Seurat, scVI and Harmony.

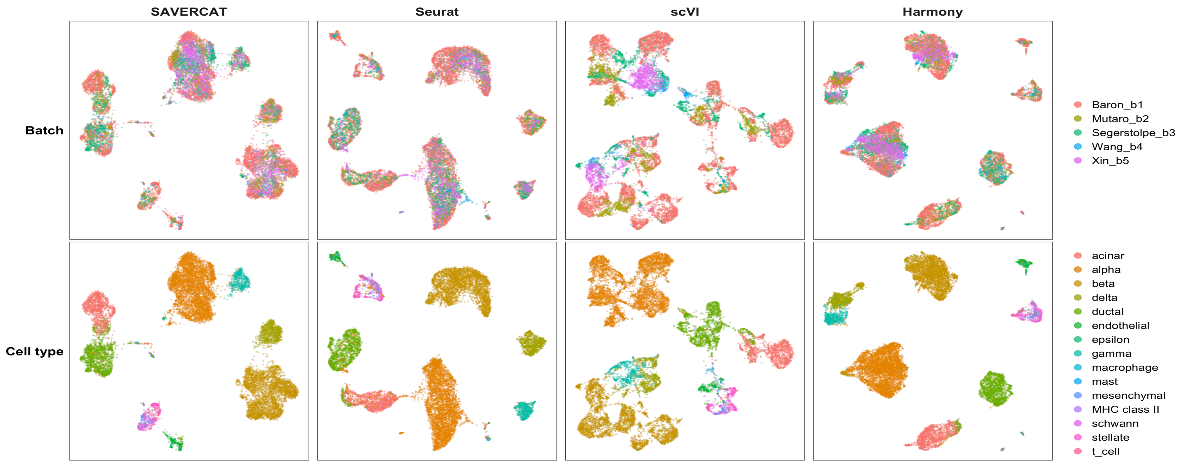


**Figure 3** UMAP visualization of 14,767 non-UMI and UMI human pancreas cells from five studies (dataset 4) to evaluate 4 methods: SAVERCAT, Seurat, scVI and Harmony.

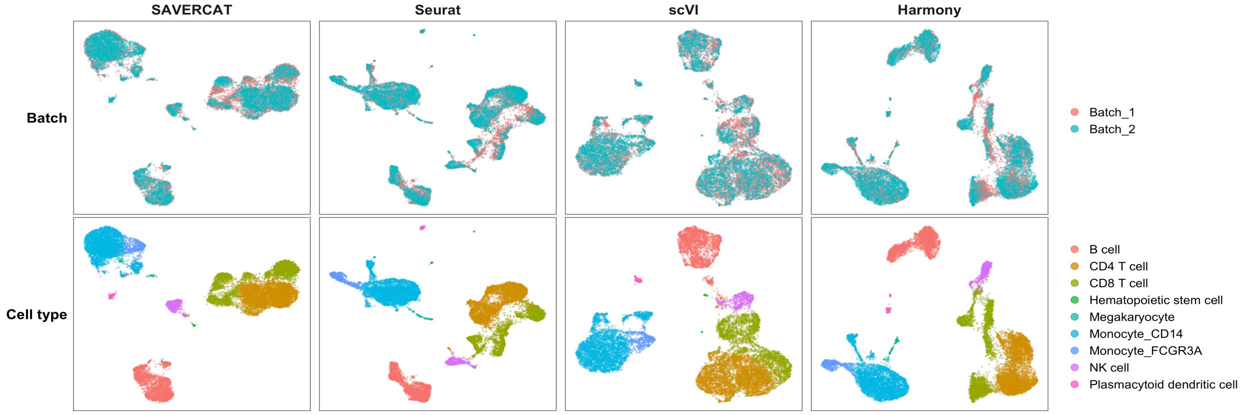


**Figure 4** UMAP visualization of 15,476 UMI human PBMC cells from two 10x Genomics technologies (dataset 5) to evaluate 4 methods: SAVERCAT, Seurat, scVI and Harmony.

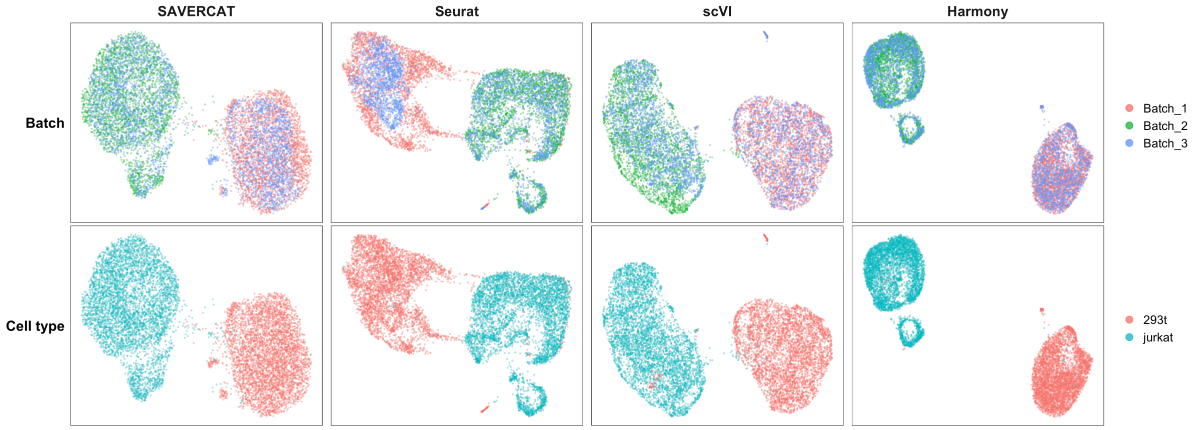


**Figure 5** UMAP visualization of 9,530 UMI human 293T and mouse 3T3 cells in three batches (dataset 6) to evaluate 4 methods: SAVERCAT, Seurat, scVI and Harmony.

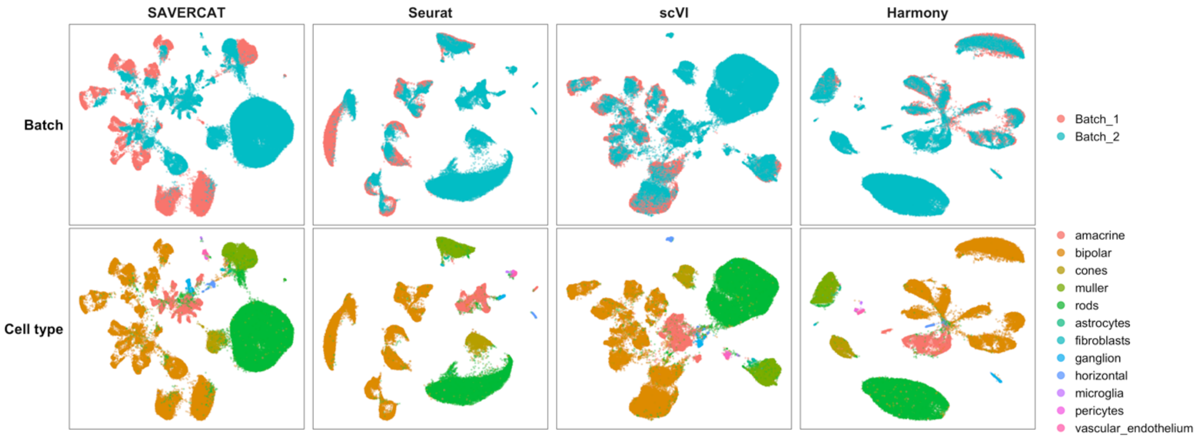


**Figure 6** UMAP visualization of 71,638 UMI mouse retina cells from two studies (dataset 7) to evaluate 4 methods: SAVERCAT, Seurat, scVI and Harmony.

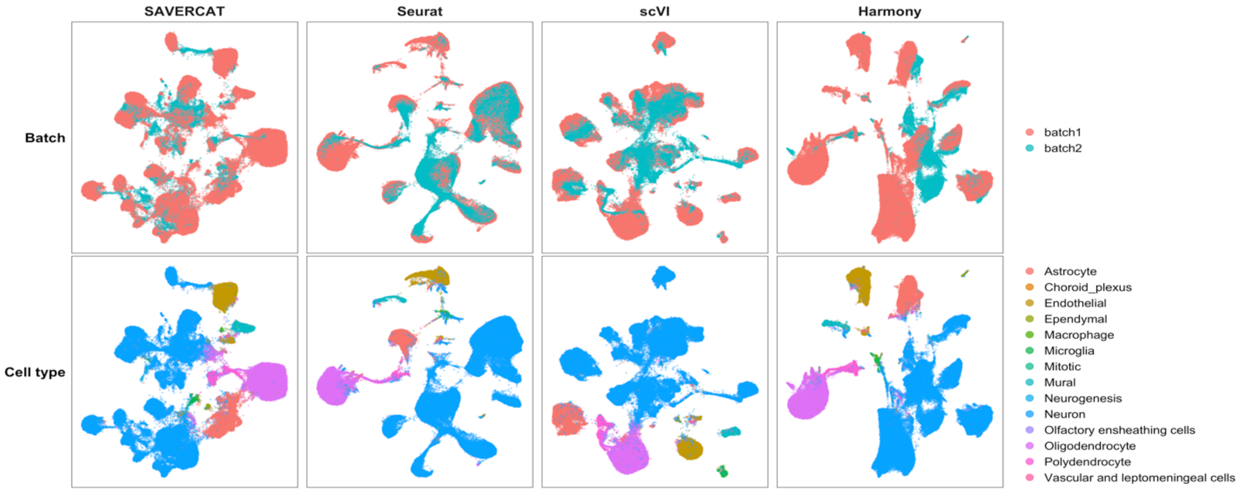


**Figure 7** UMAP visualization of 100,000 sub-sampled UMI mouse brain cells from two studies (dataset 8) to evaluate 4 methods: SAVERCAT, Seurat, scVI and Harmony.

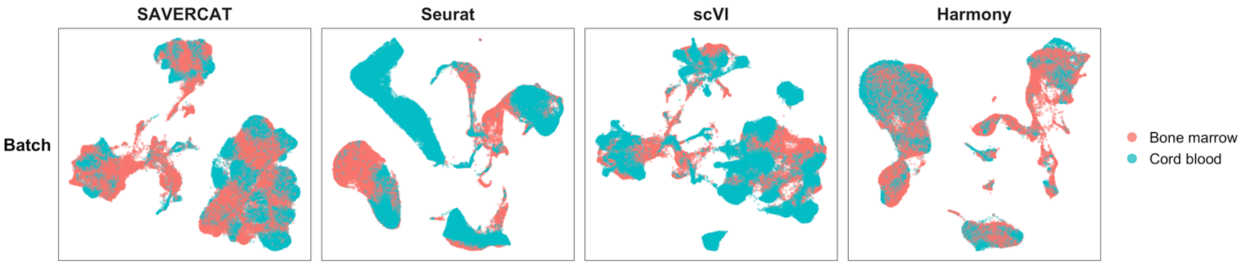


**Figure 8** UMAP visualization of 100,000 sub-sampled UMI bone marrow and cord blood-derived cells from the Human Cell Atlas (dataset 8) to evaluate 4 methods: SAVERCAT, Seurat, scVI and Harmony.

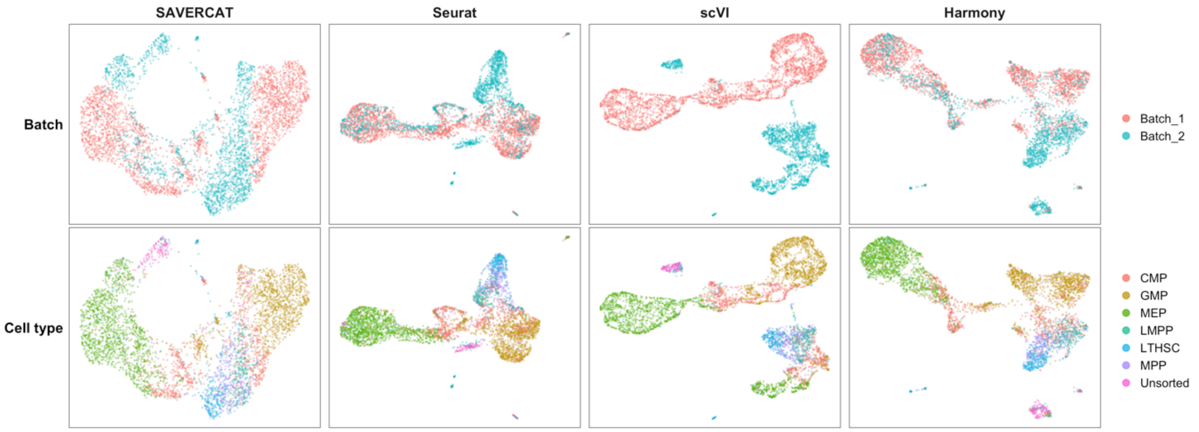


**Figure 9** UMAP visualization of 4,649 non-UMI and UMI mouse hematopoietic stem and progenitor cells from two studies (dataset 10) to evaluate 4 methods: SAVERCAT, Seurat, scVI and Harmony.

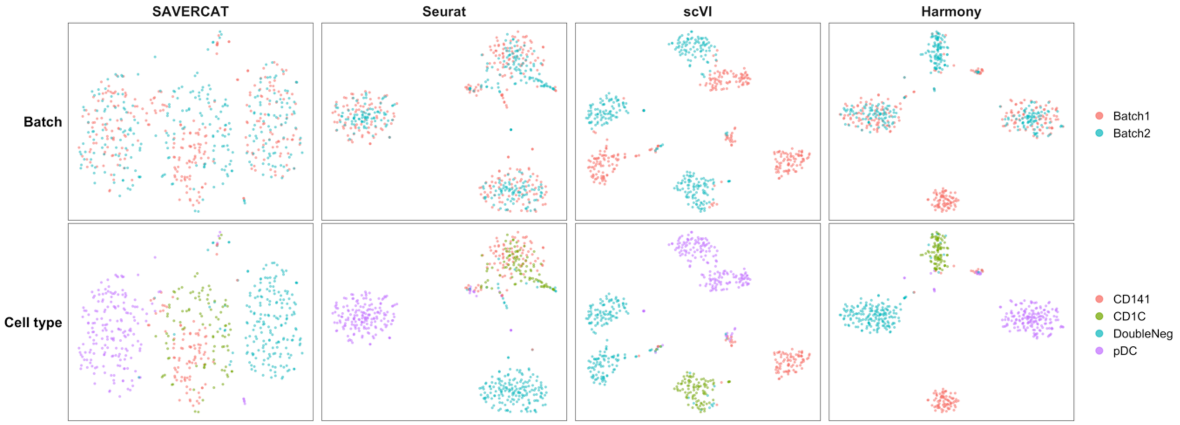


**Figure 10** t-SNE visualization of 576 non-UMI human dendritic cells from 2 batches (dataset 1) to evaluate 4 methods: SAVERCAT, Seurat, scVI and Harmony.

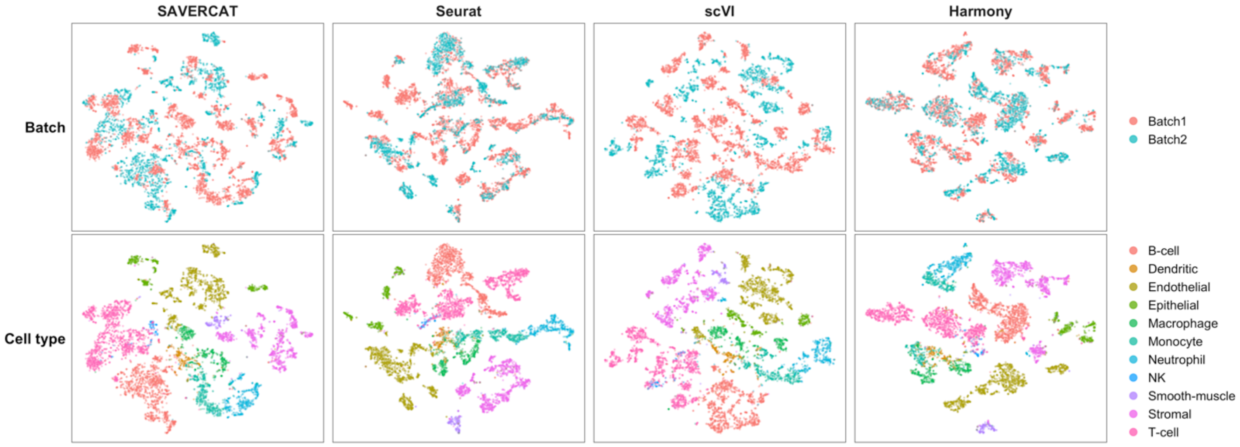


**Figure 11** t-SNE visualization of 6,954 non-UMI cells from two Mouse Cell Atlas studies (dataset 2) to evaluate 4 methods: SAVERCAT, Seurat, scVI and Harmony.

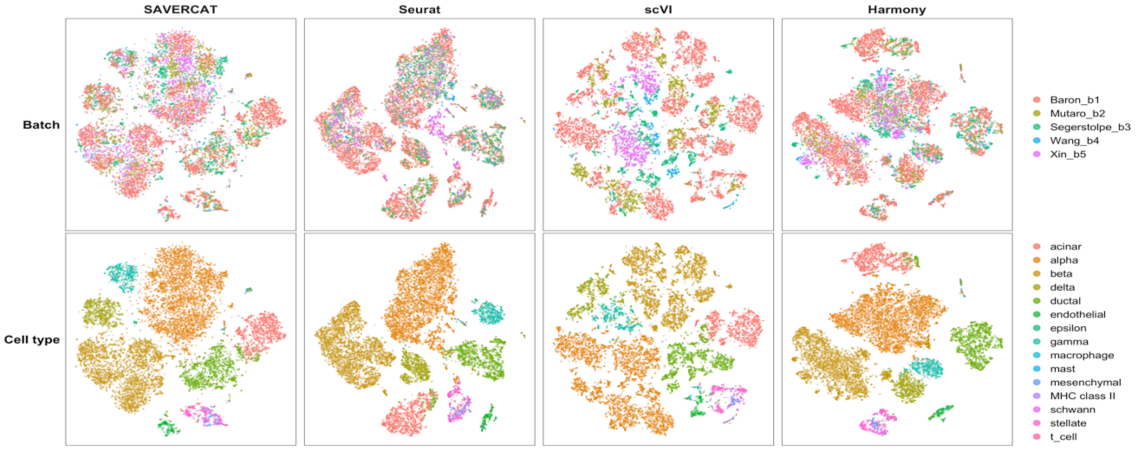


**Figure 12** t-SNE visualization of 14,767 non-UMI and UMI human pancreas cells from five studies (dataset 4) to evaluate 4 methods: SAVERCAT, Seurat, scVI and Harmony.

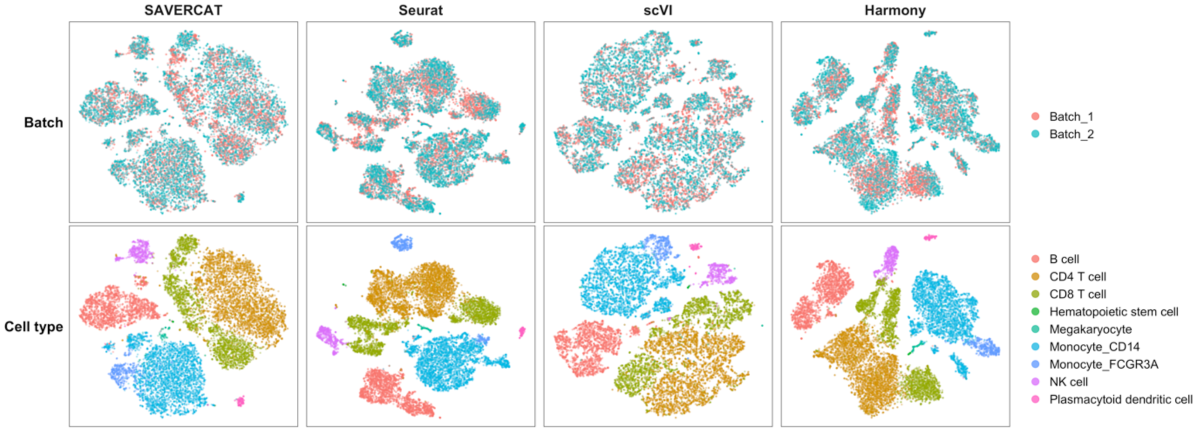


**Figure 13** t-SNE visualization of 15,476 UMI human PBMC cells from two 10x Genomics technologies (dataset 5) to evaluate 4 methods: SAVERCAT, Seurat, scVI and Harmony.

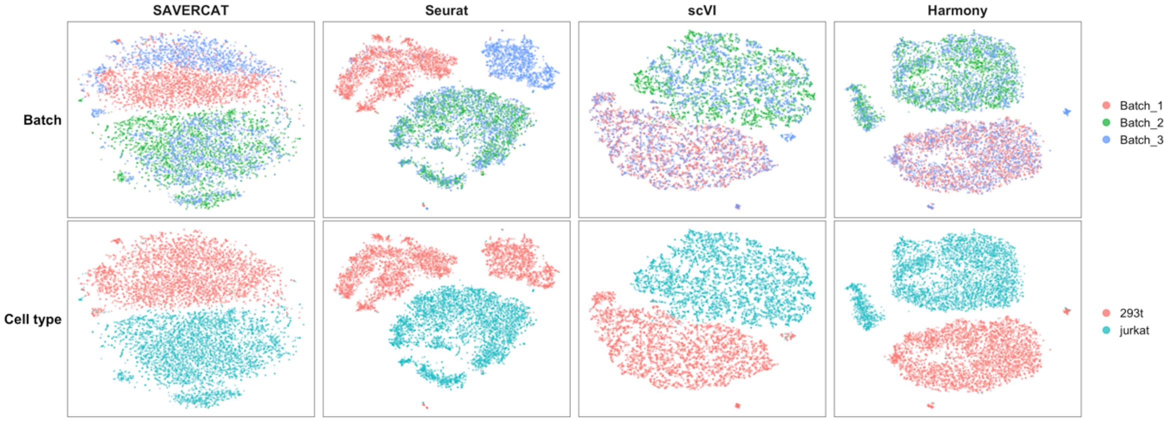


**Figure 14** t-SNE visualization of 9,530 UMI human 293T and mouse 3T3 cells in three batches (dataset 6) to evaluate 4 methods: SAVERCAT, Seurat, scVI and Harmony.

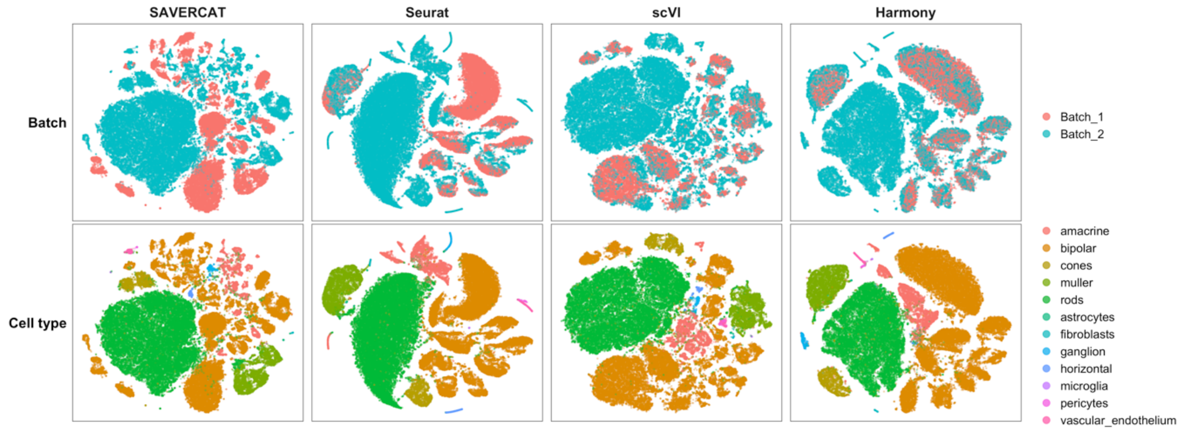


**Figure 15** t-SNE visualization of 71,638 UMI mouse retina cells from two studies (dataset 7) to evaluate 4 methods: SAVERCAT, Seurat, scVI and Harmony.

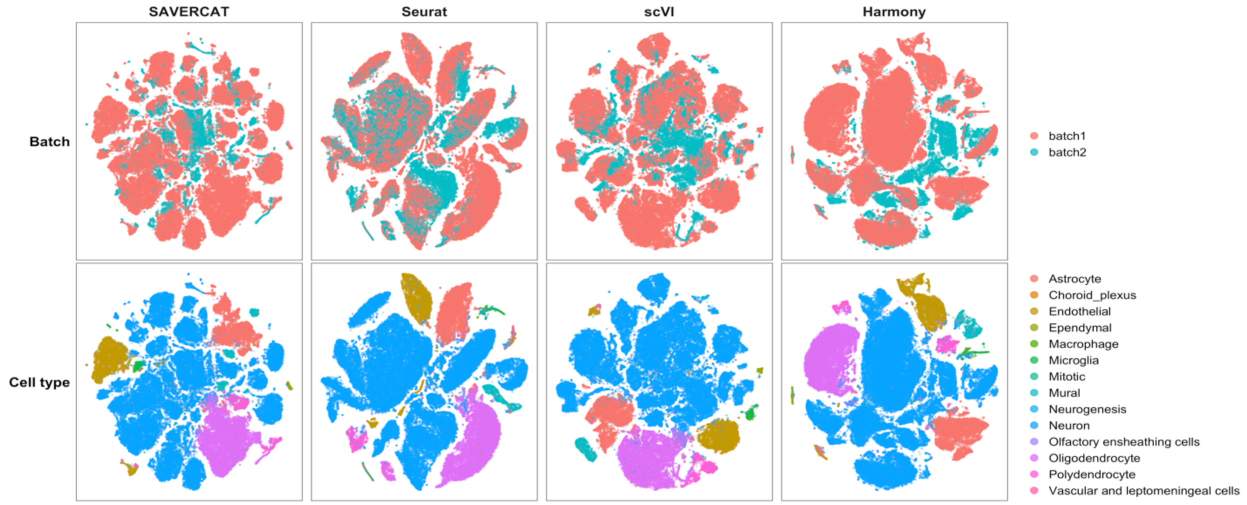


**Figure 16** t-SNE visualization of 100,000 sub-sampled UMI mouse brain cells from two studies (dataset 8) to evaluate 4 methods: SAVERCAT, Seurat, scVI and Harmony.

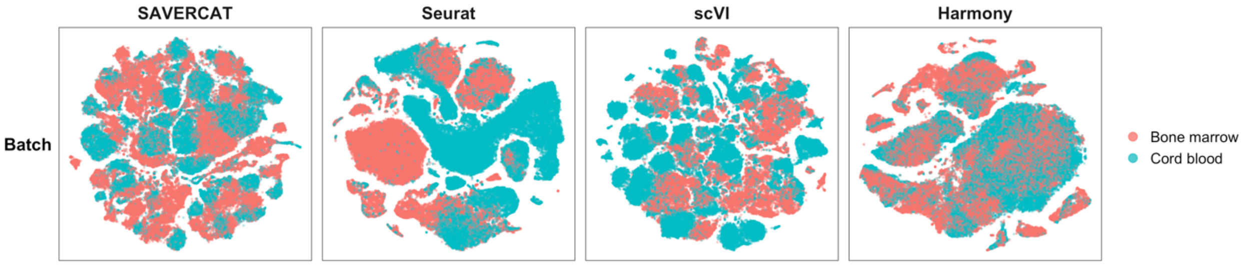


**Figure 17** t-SNE visualization of 100,000 sub-sampled UMI bone marrow and cord blood-derived cells from the Human Cell Atlas (dataset 8) to evaluate 4 methods: SAVERCAT, Seurat, scVI and Harmony.

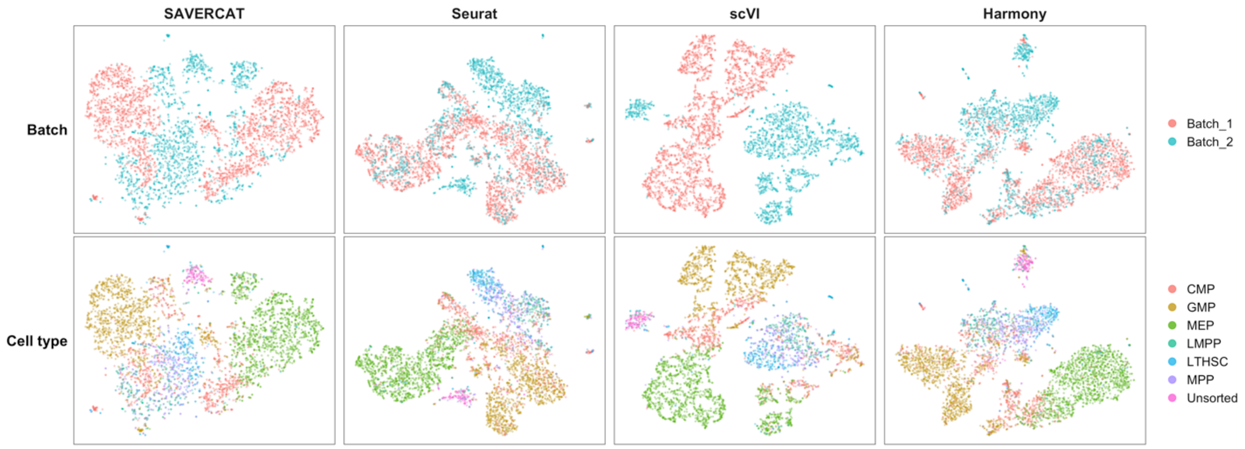


**Figure 18** t-SNE visualization of 4,649 non-UMI and UMI mouse hematopoietic stem and progenitor cells from two studies (dataset 10) to evaluate 4 methods: SAVERCAT, Seurat, scVI and Harmony.
